# Peripheral Auditory Nerve Impairment in a Mouse Model of Syndromic Autism

**DOI:** 10.1101/2022.06.02.494499

**Authors:** Nathan McChesney, Jeremy L. Barth, Jeffrey A. Rumschlag, Junying Tan, Adam J. Harrington, Kenyaria V. Noble, Carolyn M. McClaskey, Phillip Elvis, Silvia G. Vaena, Martin J. Romeo, Kelly C. Harris, Christopher W. Cowan, Hainan Lang

## Abstract

Dysfunction of the peripheral auditory nerve (AN) contributes to dynamic changes throughout the central auditory system, resulting in abnormal auditory processing, including hypersensitivity. Altered sound sensitivity is frequently observed in autism spectrum disorder (ASD), suggesting that AN deficits and changes in auditory information processing may contribute to ASD-associated symptoms, including social communication deficits and hyperacusis. The MEF2C transcription factor is associated with risk for several neurodevelopmental disorders, and mutations or deletions of *MEF2C* produce a haploinsufficiency syndrome characterized by ASD, language and cognitive deficits. A mouse model of this syndromic ASD (i.e., *Mef2c*^+/-^ or *Mef2c*-Het) recapitulates many of the *MEF2C* Haploinsufficiency syndrome-linked behaviors including communication deficits. We show here that *Mef2c*-Het mice exhibit functional impairment of the peripheral AN and a modest reduction in hearing sensitivity. We find that MEF2C is expressed during development in multiple AN and cochlear cell types, and in *Mef2c*-Het mice, we observe multiple cellular and molecular alterations associated with the AN, including abnormal myelination, neuronal degeneration, neuronal mitochondria dysfunction, and increased macrophage activation and cochlear inflammation. These results reveal the importance of MEF2C function in inner ear development and function and the engagement of immune cells and other non-neuronal cells, which suggests that microglia/macrophages and other non-neuronal cells might contribute, directly or indirectly, to AN dysfunction and ASD-related phenotypes. Finally, our study establishes a comprehensive approach for characterizing AN function at the physiological, cellular, and molecular levels in mice, which can be applied to animal models with a wide range of human auditory processing impairments.

**Significance Statement:** This is the first report of peripheral auditory nerve (AN) impairment in a mouse model of human *MEF2C* haploinsufficiency syndrome that has well-characterized ASD related behaviors including communication deficits, hyperactivity, repetitive behavior, and social deficits. We identify multiple underlying cellular, sub-cellular, and molecular abnormalities that may contribute to peripheral AN impairment. Our findings also highlight the important roles of immune cells (e.g., cochlear macrophages) and other non-neuronal elements (e.g., glial cells and cells in the stria vascularis) in auditory impairment in ASD. The methodological significance of the study is the establishment of a comprehensive approach for evaluating peripheral AN function and impact of peripheral AN deficits with minimal hearing loss.

## Introduction

Children with hearing impairment have an increased risk for developing other disabilities such as autism spectrum disorder (ASD). Hearing impairment might contribute to ASD core symptoms by interfering with language development, communication, social interaction, and overall quality of life (Kancherla et al., 2013; Do et al., 2017). Approximately 1 in 59 children with hearing loss receives school-based services for ASD, which is a much higher incidence than the 1 in 110 children with normal hearing reported to have ASD (Szymanski et al., 2012). It’s difficult to disambiguate the effects of ASD from the effects of hearing loss and communication when both are present (Garreau et al., 1984; Roper et al., 2003). Previous studies found peripheral auditory system deficits in ASD children. For example, auditory brainstem response (ABR) wave I latencies were significantly delayed in an ASD group, suggesting peripheral auditory nerve (AN) dysfunction (Rosenhall et al., 2003). ASD children also show a reduction in otoacoustic emissions at the mid-frequency region compared to age-matched controls, indicating an abnormality of outer hair cell activity (Bennetto et al., 2017).

Although previous observations support the hypothesis that central auditory processing abnormalities and structural/functional alterations contribute to ASD deficits, the direct impact of peripheral auditory system abnormalities on ASD symptoms is largely unknown (Rojas et al., 2011; Edgar et al., 2015; Rotschafer and Cramer 2017; Scott et al., 2018; Smith et al., 2019). Importantly, accumulating evidence demonstrates that peripheral auditory system deficits, e.g., dysfunction/damage of peripheral AN or inner hair cells, contribute to changes observed in higher levels of the central auditory system (Chambers et al., 2016; Salvi et al., 2017; Schrode et al., 2018; Parthasarathy et al., 2019; Auerbach et al., 2021). These changes may contribute to auditory hypersensitivity, or hyperacusis, which is defined as unusual intolerance of ordinary environmental sounds and is commonly seen in ASD patients (Hickox and Liberman 2014; Knipper et al., 2013; Chambers et al., 2016; Salvi et al., 2017; Amir et al., 2018). Given that ASD behaviors are linked to deficits at multiple levels of the nervous system (Amaral et al., 2008; Ha et al., 2015), it is challenging to distinguish the contribution of peripheral AN dysfunction to communication deficits and other ASD-like behaviors from the contributions of dysfunction at higher levels of the auditory pathway. Therefore, animal models of human ASD with a well-characterized peripheral auditory component are vitally needed to advance our understanding of the link between peripheral auditory impairment and ASD-related behaviors.

MEF2C belongs to the myocyte enhancer factor 2 (MEF2) subfamily of the MADS (MCM-agamous-deficiens-serum response factor) gene family, which is produced in several types of neurons and plays an important regulatory role in neuronal differentiation and synapse density in the developing cortex (Leifer et al., 1993; Harrington et al., 2016; Tu et al., 2017). Human *MEF2C* haploinsufficiency syndrome (MCHS), often caused by gene microdeletion or point mutation, is a newly described neurodevelopmental disorder with common symptoms that include ASD, absence of speech, hyperactivity, seizures, and intellectual disability. Studies of MCHS patients have identified numerous novel mutations in *MEF2C*, with many clustering in the DNA binding and dimerization domains and producing loss of DNA binding (Harrington et al., 2020). Based on these findings, we generated a DNA binding-deficient global *Mef2c* heterozygous mouse model of MCHS (*Mef2c*-Het mice). These mice exhibit numerous ASD-like behaviors, such as reduced ultrasonic vocalizations in a social context, social interaction deficits, hyperactivity, repetitive behavior, and reduced sensitivity to painful stimuli (Harrington et al., 2020). This mouse model of syndromic autism provides a means to directly investigate the relationship between peripheral AN dysfunction and ASD behaviors. Here we report that this mouse model of human MCHS has functional impairments of the AN at the physiological, cellular, and molecular levels. Further, our findings highlight the importance of immune cells and non-neuronal elements as regulatory factors in auditory impairment in ASD.

## Materials and Methods

### Animals

All aspects of animal research were conducted in accordance with the guidelines of the Institutional Animal Care and Use Committee of the Medical University of South Carolina (MUSC). CBA/CaJ mice, originally purchased from the Jackson Laboratory (Bar Harbor, ME), were bred in a low-noise environment at the Animal Research Facility at MUSC. *Mef2c*^*+/−*^ (*Mef2c*-Het) mice were generated previously as described (Harington et al., 2020). Briefly, this global heterozygous *Mef2c* mutant line was established by crossing exon 2 floxed *Mef2c* mice with *Prm1*-Cre mice (#003328; The Jackson Laboratory, Bar Harbor, ME) to create a constitutive *Mef2c* loss-of-function allele (*Mef2c*^+/Δ exon2^). The *Prm1*-Cre transgene was subsequently removed during repeated backcrossing to C57BL/6J wild-type mice. This global heterozygous mutant impacts the Mef2c DNA binding function, but not the protein expression. In *Mef2c* Het mice, the germline-transmitted *Mef2c* heterozygous loss-of-function allele was generated by crossing Prm1-Cre (sperm expression) males with females having floxed *Mef2c* exon 2, which encodes a portion of the MADS/MEF2 domains. Removal of exon 2 allows for in-frame splicing of exon 1 to 3, producing a near full-length *Mef2c* open reading frame that lacks only the portion encoding the DNA binding/dimerization domain, similar to the MCHS missense mutations observed in patients. *Mef2c* conditional heterozygous mice were generated by crossing *Mef2c-flox* mice with macrophage-selective Cre-expressing transgenic mice (*Cx3Cr1*^*creER/creER*^; Jackson Laboratory #021160) to generate *Mef2c*^*fl/+*^; Cx3cr1-*Cre* mice (*Mef2c cHet*^*Cx3cr1*^ mice) that were compared to their flox-negative littermates (control mice).

All mice received food and water *ad libitum* and were maintained on a 12-hour light/dark cycle. Mice with signs of obstruction or infection in the external ear canal or middle ear were excluded. All animals underwent measurement of auditory physiology and at least one other molecular or cellular assay, including RNA sequencing, immunohistochemistry, and transmission electron microscopy (see detailed information of these assays below).

### Auditory Physiology

Cochlear and auditory nerve function was measured using the cochlear microphonic (CM), and the auditory brainstem response (ABR). The CM is used as a proxy of the mouse cochlear health (Cheatham et al., 2011). The CM is an AC receptor potential with electromechanical origins that is sensitive to changes in the endocochlear potential (EP) and hair cell (HC) loss. The CM represents the functional sum of the hair cells, basilar membrane, and strial effects. The CM is often considered a measure of OHC function, similar to OAEs, such that damage to the OHCs and hearing loss result in reduced CM amplitude. The ABR wave I was used to assess several metrics of the peripheral AN function via both averaged and single-trial ABR recording (detailed information in sections below). Averaged recording provided estimates of ABR wave I threshold and suprathreshold measurement and single-trial ABR recordings provided estimates of neural synchrony as in our previous reports in mouse models (Jyothi et al., 2010; McClaskey et al., 2020; Panganiban et al., 2018; 2021). Suprathreshold estimates of AN peak response amplitudes and latency, and neural synchrony (Phase-locking value: PLV) were acquired. These suprathreshold measures of the ABR have been associated with speech recognition in noise (Harris et al., 2018, 2021).

For measurements of auditory function, wildtype (WT) and *Mef2c*-Het littermates (both male and female) at postnatal day (P) 16, and 2-3 months were anesthetized via an intraperitoneal injection of a cocktail containing 20 mg/kg xylazine and 100 mg/kg ketamine. Auditory tests were performed in a sound-isolation booth. CM and ABR recordings were performed using a TDT RZ6 multi-I/O. Equipment for auditory functional measurements (CM and ABR) was calibrated before use with TDT RPvdsEx software (Tucker Davis Technologies, Gainsville, FL) and a model 378C01 ICP microphone system provided by PCB Piezotronics, Inc. (Depew, NY). In a closed-field setup, sound stimuli were delivered into the ear canal via a 3–5 mm diameter tube.

#### Cochlear Microphonic

The CM was elicited by a 100-ms 4 kHz tone-burst with a 10 ms cos^2^ rise/fall time. Sound levels were reduced in 10 dB steps, from 120 to 70 dB SPL. Since the CM reflects a presynaptic potential, stimuli had a high repetition rate (5 ms gap between presentations) to reduce the influence of the neural response. The active electrode (channel1) was placed was placed around the bulla of the ear for which the CM was measured and the inverting electrode into the scalp subdermally at the vertex and between the ears. The ground electrode was placed at the hindlimb. The CM was recorded using 600 trials. Data processing was performed in MATLAB using custom scripts and EEGLAB. The CM was measured in the frequency domain as the power of the signal at the eliciting stimulus frequency (4 kHz).

#### ABR: Threshold measures

ABRs were evoked at frequencies of 4, 5.6, 11.3, 16, 22.6, 32, 40, and 45.2 kHz with 5-ms tone pips with 0.5 ms cos^2^ rise/fall times delivered at 31 times/s using a TDT system III with a RP2.1 enhanced real-time processor. For ABR recordings, the active electrode (channel1) was placed was placed into the scalp subdermally at the vertex and between the ears, and the inverting electrode around the bulla of the ear for which the ABR was measured. The ground electrode was placed at the hindlimb. Sound levels were reduced by 5 dB steps, from 90 to 10 dB SPL. The average ABR waveform for each level tested consisted of 256 trials. Wave I thresholds were determined visually for each mouse as the lowest level that elicited an ABR response, as determined by at least two experienced reviewers. Thresholds were then averaged at each frequency across groups, and the mean ± standard error of the mean was calculated and plotted using Origin 6.0 software (OriginLab Corporation, Northampton, MA) or GraphPad Prism 9 (GraphPad Software, San Diego, CA).

#### ABR: Suprathreshold measures

Suprathreshold wave I peak amplitudes were measured. Peak-to-peak wave I amplitudes were measured from the positive peak to the following negative deflection using a custom MATLAB (MathWorks, Inc., Natick, MA) script, in which the experimenter selected peaks while being blinded to genotype. Our recent report (McClaskey et al., 2020) identified a breakpoint in the growth of wave I at around 75 dB SPL. Therefore, levels above this breakpoint were classified as suprathreshold and reflected the response of a larger contingent of AN fibers with various thresholds, including higher threshold fibers hypothesized to be important for speech recognition.

The procedure for continuous single-trial recording of ABR was modified from our recent report (McClaskey et al., 2020; Panganiban et al., 2021). Stimuli were 1.1 ms in duration with 0.55-ms cos^2^ rise/fall times and were presented at a rate of 21 /s at the levels of 5-90 dB SPL in 5 dB increments. At least 500 tone pips of each sound level were presented. ABR responses were stored and processed offline in MATLAB using the EEGlab toolbox (Delorme & Makeig, 2004) and the ERPlab extension (Lopez-Calderon & Luck, 2014). Continuous data were bandpass filtered between 100 Hz and 3000 Hz using an 8^th^ order Butterworth filter. Data were then epoched from -3 to +11 ms relative to stimulus onset and baseline corrected by subtracting the mean value in the pre-stimulus window of -3 to -1 ms. Epochs with voltages exceeding +/-5 µV were rejected and the remaining epochs were visually inspected for excessive movement or other artifacts. Artifact-free trials were saved in the single-trial form in EEGLAB (https://sccn.ucsd.edu/eeglab/index.php) and as a trial-averaged ABR waveform in ERPLAB (https://sccn.ucsd.edu/eeglab/index.php). Wave I was visually identified as the first major positive deflection following stimulus onset. Phase locking value (PLV) was calculated from the single-trial-level data.

PLV, also known as inter-trial coherence (ITC), is the length of the vector that is formed by averaging the complex phase angles of each trial at each frequency and was obtained via time-frequency decomposition of the single-trial-level data. With increasing stimulus level, first-spike latencies decrease and become more consistent across AN fibers (Heil and Irvine 1997; Heil 2004), this decrease in jitter is hypothesized to improve timing across trials, resulting in stronger PLV. Time-frequency decomposition was performed with Hanning FFT tapers via EEGlab’s newtimef() function, using 16 linearly spaced frequencies from 190 to 3150 Hz. PLV at each time point and each frequency was then calculated by taking the absolute value of the complex inter-trial coherence output of the newtimef() function, using the following equation, in which *F*_*k*_ (f, t) is the spectral estimate of trial *k* at frequency *f* and time *t*:

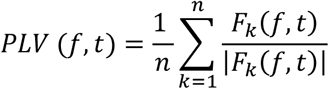

The maximum PLV between 250 and 2500 Hz in the 2-ms window surrounding the wave I peak was then measured as PLV.

### AN tissue collection and total RNA isolation

Note that throughout the paper, the term “auditory nerve” (AN) refers to the peripheral part of the VIII nerve extending from synapses with hair cells in the organ of Corti to the main nerve trunk within the modiolus and extending into the internal auditory canal. AN tissues were collected at their designated end-points using both cochleae from each mouse. Microdissections were performed to isolate the AN from the rest of the cochlear structures, taking care to preserve peripheral fibers. Samples were created by pooling ANs collected from either one mouse (aging study) or two mice (Mef2c deficiency study). Total RNA was isolated using the RNeasy Plus Mini Kit (Qiagen Inc, Germantown, MD) per the manufacturer’s instructions. Quality of each total RNA preparation was assessed by 4200 TapeStation (Agilent Technologies, Santa Clara, CA). Low-quality samples showing degradation or contamination were excluded.

### RNA sequencing (RNA-seq)

RNA-seq analysis of CBA/CaJ mouse AN development (NCBI accession GSE133823) and aging (NCBI accession GSE141865) have been described previously (Panganiban et al., 2021). For the developmental study, P3, P7, P14, P21 and adult AN samples were analyzed; for the aging study, aged adult (2.5 years) and young adult (2 months) AN samples were analyzed. RNA-seq analysis of the effects of Mef2c deficiency was done using P21 WT and *Mef2c*-Het mice (both sexes) and four biological replicates of each sample type. Sequencing libraries were prepared at the Medical University of South Carolina Translational Science Lab. Polyadenylated RNA was captured from 50 ng of total RNA per sample and libraries were prepared using the NEBNext^®^ Poly(A) mRNA Magnetic Isolation Module and NEBNext^®^ Ultra^™^ II Directional RNA Library Prep Kit for Illumina^®^ (New England Biolabs, Ipswich, MA). Paired-end sequencing was done at the Vanderbilt VANTAGE core laboratory (Vanderbilt University, Nashville, TN) to a depth of 25 million reads/library using an Illumina NovaSeq 6000. Sequencing data were analyzed using Partek® Flow® software. Reads were aligned to mouse genome assembly mm10 with an implementation of STAR (Spliced Transcripts Alignment to a Reference) (Dobin et al., 2013) and quantified to annotation model (Partek E/M) using mm10. Calculation of fold change and adjusted p-value (FDR step up) was done with DESeq2 (Love et al., 2014). Raw sequencing data (fastq files) and comparison results for the *Mef2c*-Het study are archived in NCBI Gene Expression Omnibus (accession GSE182857).

### Comparative analysis of RNA-seq data

For evaluation of ASD-related gene expression during AN development: 1) comparison data were obtained for GSE133823; and 2) ASD-related genes were compiled from SFARI database categories 1-4, which comprise genes with high confidence through genes with minimal evidence (https://gene.sfari.org). Genes differentially expressed during development were defined as adjusted p-value (FDR step up) <0.10 and absolute fold change >1.5 for P21 versus P3 AN. Commonality between developmentally regulated genes, ASD-related genes, and macrophage/inflammation-related genes was done by Venn analysis using gene symbols as input. For RNA-seq analysis of *Mef2c*-Het compared to WT AN, differential expression was also defined as adjusted p-value (FDR step up) <0.10 and absolute fold change >1.5, identifying 258 genes. Biological process enrichment was conducted with ToppFun (Chen et al., 2009); enrichment results were refined and the interactive graph generated by REVIGO (Supek et al., 2011). To determine if genes affected by Mef2c deficiency were also affected in aged AN, comparison data were obtained for the aging AN RNA-seq study (accession GSE141865) and statistical measures for the 258 genes affected in *Mef2c*-Het AN were reviewed. Significant difference for aged adult versus young adult AN was defined as adjusted p-value (FDR step up) <0.10, identifying 19 of the 258 genes.

### Immunohistochemistry and Lectin histochemistry

Immunohistochemical procedures were modified from previous studies (Panganiban et al., 2018; 2021). After end-point physiological recordings, mouse cochleae were collected and immediately fixed with 4% paraformaldehyde solution in 1x phosphate-buffered saline for 2 hours at room temperature and decalcified with 0.12 M ethylenediamine tetraacetic acid (EDTA) at room temperature for 1-2 days. To prepare mouse cochlea sections, cochleae were embedded in the Tissue-Tek OCT compound and sectioned at a thickness of approximately 12 μm. For whole-mount preparations of mouse cochlear tissues, the ANs, cochlear lateral wall, and organs of Corti were isolated separately based on the experimental plans (e.g., to count hair cells, or to evaluate the strial microvasculature).

Primary and secondary antibodies used for immunohistochemistry are listed in **Table 1**. Staining was performed by either indirect method using biotinylated secondary antibodies conjugated with fluorescent avidin (Vector Labs, Burlingame, CA), or direct method using primary antibodies conjugated to Alexa Fluor Dyes (ThermoFisher Scientific, Waltham, MA). Nuclei were counterstained using propidium iodide (PI), or 4′,6-diamidino-2-phenylindole (DAPI).

**Table 1.**
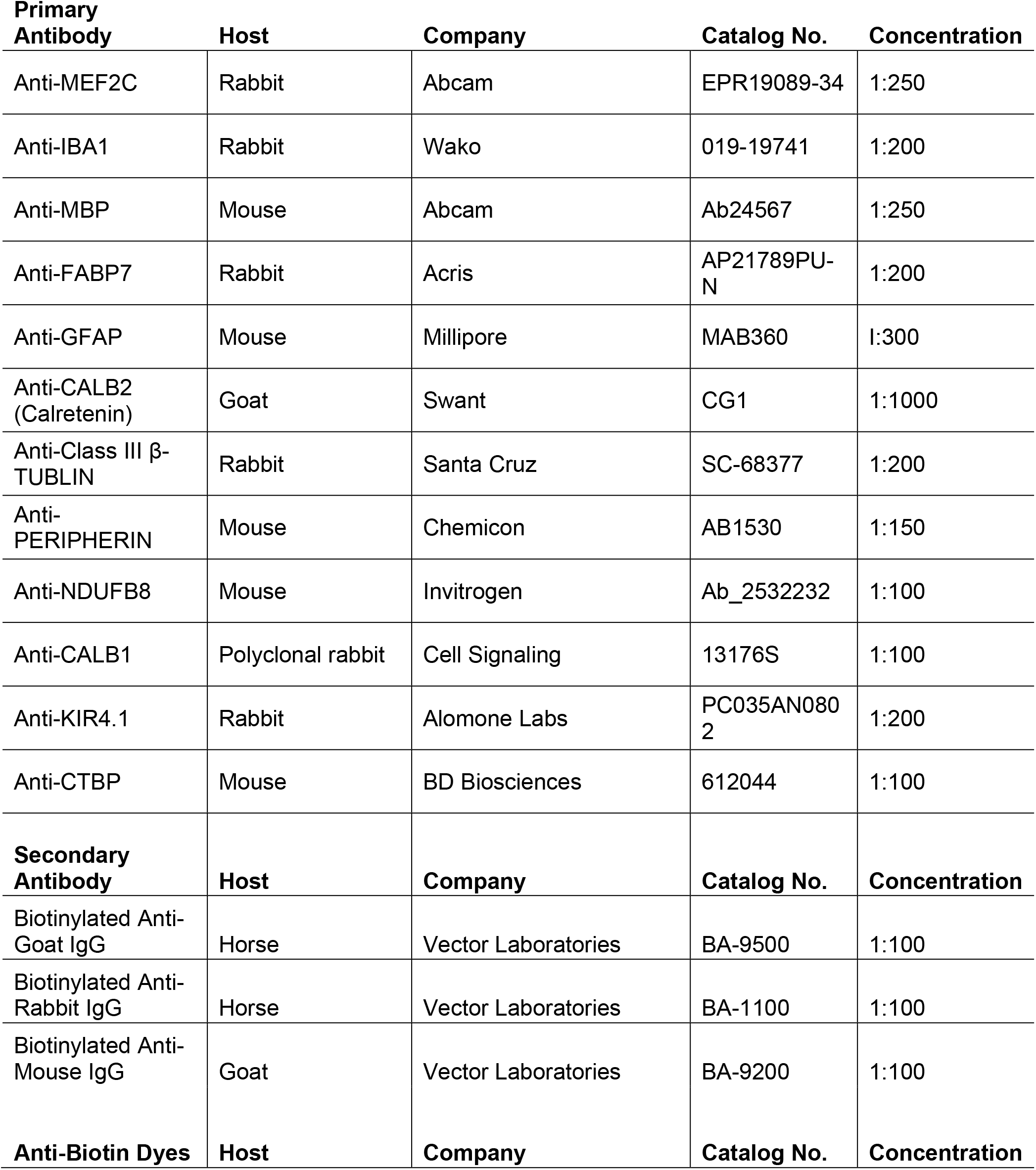
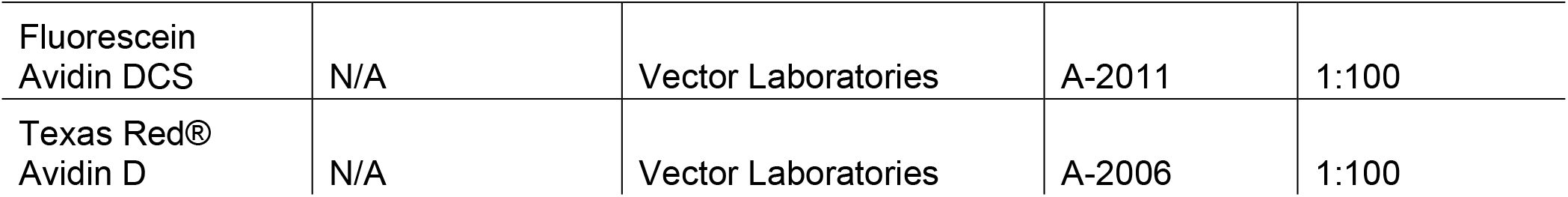
Antibodies used in the study.

| **Primary Antibody** | **Host** | **Company** | **Catalog No.** | **Concentration** |
| --- | --- | --- | --- | --- |
| Anti-MEF2C | Rabbit | Abcam | EPR19089-34 | 1:250 |
| Anti-IBA1 | Rabbit | Wako | 019-19741 | 1:200 |
| Anti-MBP | Mouse | Abcam | Ab24567 | 1:250 |
| Anti-FABP7 | Rabbit | Acris | AP21789PU-N | 1:200 |
| Anti-GFAP | Mouse | Millipore | MAB360 | 1:300 |
| Anti-CALB2 (Calretenin) | Goat | Swant | CG1 | 1:1000 |
| Anti-Class III β-TUBLIN | Rabbit | Santa Cruz | SC-68377 | 1:200 |
| Anti-PERIPHERIN | Mouse | Chemicon | AB1530 | 1:150 |
| Anti-NDUFB8 | Mouse | Invitrogen | Ab_2532232 | 1:100 |
| Anti-CALB1 | Polyclonal rabbit | Cell Signaling | 13176S | 1:100 |
| Anti-KIR4.1 | Rabbit | Alomone Labs | PC035AN0802 | 1:200 |
| Anti-CTBP | Mouse | BD Biosciences | 612044 | 1:100 |
| **Secondary Antibody** | **Host** | **Company** | **Catalog No.** | **Concentration** |
| Biotinylated Anti-Goat IgG | Horse | Vector Laboratories | BA-9500 | 1:100 |
| Biotinylated Anti-Rabbit IgG | Horse | Vector Laboratories | BA-1100 | 1:100 |
| Biotinylated Anti-Mouse IgG | Goat | Vector Laboratories | BA-9200 | 1:100 |
| **Anti-Biotin Dyes** | **Host** | **Company** | **Catalog No.** | **Concentration** |
| Fluorescein Avidin DCS | N/A | Vector Laboratories | A-2011 | 1:100 |
| Texas Red® Avidin D | N/A | Vector Laboratories | A-2006 | 1:100 |

A lectin histochemical approach was used to visualize the blood vessel system in mouse strial vascularis modified from previous studies (Meyer et al., 2008; Shi 2011). For whole-mount preparations of mouse cochlear lateral wall tissues, the fluorescent lectin Gs-IB4 Alexa Fluor® 488 was diluted in PBS with 0.1% CaCl_2_ (1:100) and the tissue was incubated overnight at 4°C.

Slice and confocal image stacks were collected using a ZEISS LSM 880 NLO with Airyscan and ZEN acquisition software (ZEISS United States, Thornwood, NY). Images were processed using ZEN 2012 Blue Edition (Carl ZEISS Microscopy GmbH) and Adobe Photoshop CC (Adobe Systems Incorporated, San Jose, CA). Quantitative analysis of macrophage size and volume was done with the Surface module of IMARIS (IMARISx64 9.3.1) with Z-stack images collected via LSM 880 NLO.

### Transmission electron microscopy (TEM)

Samples were prepared for TEM using procedures modified from previous publications (Lang et al., 2015). Briefly, deeply anesthetized mice were cardiac perfused with 15 mL of a fixative solution containing 4% paraformaldehyde and 2% glutaraldehyde in 0.1 M phosphate buffer, pH 7.4). The same fixative solution was used to perfuse the excised cochleae through the round window and for further immersion overnight at 4 °C. Cochleae were decalcified using 0.12 M EDTA solution at room temperature for 2–3 days with a magnetic stirrer. Cochleae were then fixed using a solution containing 1% osmium tetroxide and 1.5% ferrocyanide for 2 hours in the dark and then dehydrated and embedded in Epon LX 112 resin. Semi-thin sections for pre-TEM observation of AN orientation were cut at 1 μm thickness and stained with toluidine blue. Once a coronal plane for a given cochlear turn was seen, ultra-thin sections at 70 nm thickness were cut and stained with uranyl acetate and lead citrate. Ultra-thin sections were examined using a JEOL JEM-1010 transmission electron microscope (JEOL USA, Inc., Peabody, MA).

### Characterization of spiral ganglion neuron pathology, myelination and mitochondrial abnormity in AN

Quantitative analyses of ultrastructural alteration in myelin and mitochondria and examination of aging-like neuronal pathology were conducted on TEM graphs collected from the middle portions of cochleas. A myelin sheath was classified as abnormal if it exhibited at least two areas with degenerative myelin features. Such features included splitting, folded or collapsed lamella, or balloon-like features (Xing et al., 2012; Helvacioglu et al., 2019). A mitochondria was classified as dysfunctional if it exhibited disrupted cristae or loss of condensed cristae (Perkins et al., 2020; Joubert and Puff, 2021; **Fig. 5M**). A spiral ganglion neuron (SGN) cell body was classified as exhibiting an aging-like feature if it contained a cluster of at least two lipofuscin-accumulated granules (aging pigment; **Fig. 5F**) (Sulzer et al., 2008; Vila 2019). For quantitative analysis of abnormal myelin and aging-like neuronal pathology, four animals per genotype and 17 to 90 neurons per animal were examined. For quantitative analysis of dysfunctional mitochondria, 3 WT and 4 *Mef2c*-Het mice were analysed and 62 to 249 mitochondria per AN were examined in randomly selected AN axons.

### Experimental design and statistical analyses

Experimental plans were designed to achieve a statistical power of ≥80% and a significance level of ≤0.05 (de Aguilar-Nascimento et al., 2005). Sample sizes (n) are indicated in each figure legend or text in the Results section. Sample size for physiological measurements (e.g., ABR wave I threshold, wave I amplitude and latency) was estimated based on effect sizes obtained from observed means and standard deviations of preliminary data and from previous studies by our laboratory and our collaborators (Lang et al., 2015; Xing et al., 2012; Panganiban et al., 2018; 2022; McClaskey et al., 2020; Harrington et al., 2020). For RNA-seq, prior analyses of mouse AN demonstrated that a sample size of n = 3 was sufficient for robust detection of genes differentially expressed with an effect size of 1.5-fold and false discovery rate less than 10% (Lang et al., 2016; Panganiban et al., 2021). For datasets other than RNA-seq, data distribution was tested using either Shapiro-Wilk or Kolmogorov-Smirnov tests for normality. The appropriate parametric or non-parametric tests were then used for the AN functional and structural data. Statistical software and packages used in this study included DeSeq2 (Love et al., 2014), Microsoft Excel, R version 3.5.2 (R Foundation for Statistical Computing, 2019), and GraphPad Prism 8 (GraphPad Software, Inc., La Jolla, CA). Type III ANOVA was used for analysis of ribbon synapse count in the auditory nerve. Student’s t test and/or linear mixed-effects regression (LMER) models were used for analyses of ABR wave I threshold, amplitude growth functions, PLV and CM. Linear regression models (LM) were used for analyses of the slope of ABR wave I amplitude growth functions. LMER and LM analyses are performed in R using the lme4 package (Bates et al., 2015). For the Mann-Whitney U test, which was applied to the analyses of the averaged cellular or subcellular structures using ultrastructural, and immunohistochemical evaluation, a *p*-value of ≤ 0.05 was considered significant. For analyses of the cellular area and volume in the cochlear macrophages, unpaired, two tailed t test with Welch’s corrections was used. For differential expression analyses of complete RNA-seq datasets, a p-adjusted (FDR step-up) value of ≤ 0.10 and absolute fold change >1.5 was considered significant; for analyses that examined subsets of RNA-seq data (<1000 genes), p-adjusted (FDR step-up) value of ≤ 0.10 was considered significant. Detailed information for statistical analyses is included in supplementary data Tables designed as **Extended data Figs 1-1, 1-2, 2-1, 3-1, 3-2, 4-1, 5-1, 6-1, 6-4**, and **7-1** (these data tables were named following the Figures to which these tables are related).

## Results

### Numerous ASD risk genes including *Mef2c* are differentially expressed during mouse peripheral AN development

The ASD risk gene *MEF2C* is highly expressed in several areas of the early developing brain (e.g., in differentiated forebrain neurons within the neocortex and dentate gyrus) and maintains its expression in multiple types of post-mitotic neurons such as glutamatergic neurons, GABAergic neurons, and Purkinje cells (Leifer et al., 1993, 1997; Lyons et al., 1995; Speliotes et al., 1996; Harrington et al., 2016). We hypothesized that *Mef2c*, and possibly other ASD risk genes, may play a role in peripheral AN development. To test this hypothesis, we performed an analysis to determine if ASD risk genes were regulated during AN development. A list of ASD risk genes was obtained from the Simons Foundation Autism Research Initiative (SFARI) database; (Categories 1-4; https://gene.sfari.org) and an RNA-seq dataset profiling gene expression during AN development was obtained (NCBI GEO accession GSE133823). Evaluation of expression data for the 671 ASD risk genes found that 319 (48%) were differentially expressed during AN development (**Fig. 1A**). Similar to observations in the central nervous system, *Mef2c* was highly expressed in the mouse AN in early postnatal development periods (e.g., P3), but then it undergoes downregulation during hearing onset. It then maintains a measurable level from P21 to adulthood (**Fig 1B, C**). Interestingly, among differentially expressed genes, *Pik3cg* is a *Mef2c* gene target and plays a regulatory role in plasticity of nucleus accumbens in a cocaine-exposed animal model (Pulipparacharuvil et al., 2008).

**Figure 1.**
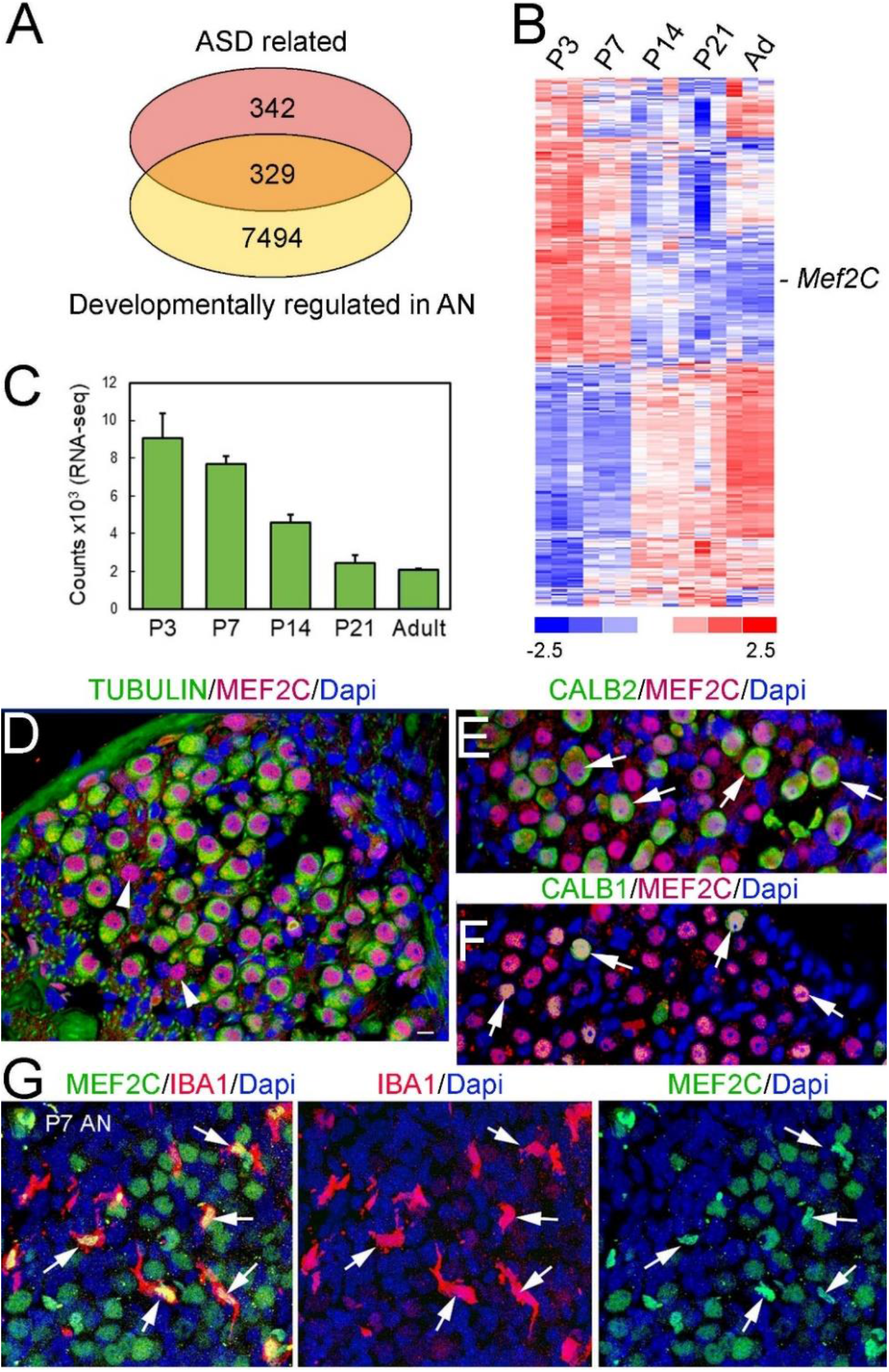
Expression of ASD risk genes in the mouse peripheral auditory system. ***A***, Venn diagram depicting overlap between 1) ASD-risk genes, and 2) genes regulated during AN development. The list of ASD-related genes differentially expressed in mouse AN during development is included in **Extended data Fig. 1-1. *B***, Heatmap of ASD-risk genes differentially expressed in mouse AN during development, including *Mef2c*. ***C***, Mef2c mRNA expression is highest in AN in postnatal stages preceding hearing onset (around P12). Graph depicts RNA-seq standardized counts (detailed information in **Extended data Fig. 1-2**). ***D-E***, Detection of MEF2C protein (red) in type SGNs (TUBULIN^+^; green in **D**), high-SR SGNs (CALB2^+^, green in **E**) and mid-SR SGNs (CALB1^+^, green in **F**) in young adult mouse AN. ***G***, Expression of MEF2C (green) in macrophages in mouse P7 AN. Macrophages were identified by immunostaining for Iba1 (red). Nuclei were stained with Dapi (blue). **Extended data Fig. 1-3 (Images)** showing MEF2C expressing cells present around a blood vessel in AN and cochlear lateral wall and areas near bone-forming cells in postnatal developing cochlea. Scale bar = 10 µm in **D** (applies to **E-G**).

To validate the *Mef2c* expression findings, immunostaining was performed for MEF2C together with markers of SGNs and macrophage/microglia, revealing that multiple cell types in AN and cochlea expressed MEF2C protein (**Fig. 1D-G**). These cell types include type I SGNs (**Fig. 1D**) (Sekerková et al., 2008; Jyothi et al., 2010), high-and mid-spontaneous rate (SR) SGNs (CALB2^+^ and CALB1^+^, respectively; **Fig. 1E,F**) (Petitpré et al., 2018; Shrestha et al., 2018; Sun et al., 2018), and macrophages (IBA1^+^, **Fig.1G**) (Brown et al., 2017). Low SR-SGN express either none or a very limited level of two calcium-binding proteins, calbindin (CALB1) and Calretinin (CALB2), which are highly expressed in mid-and high-SR fibers, respectively. We observed some MEF2Cpositive SGNs that were negative for CALB2 and CALB1 (red), suggesting that low-SR SGNs are also MEF2C-expressing neurons (**Fig. 1E,F**). MEF2C-expressing cells were also seen in and around CD31^+^ blood vessels within the strial vascularis and AN (**Extended data Fig. 1-3B,C,F,G**) (Lang et al., 2016). In addition, MEF2C was highly expressed in bone-forming cells around the cochlear lateral wall and AN (**Extended data Fig. 1-3A,E**).

### Mef2c deficiency results in a mild reduction of hearing sensitivity in young adult animals

ABR wave I thresholds were measured at P16 and in young adult (2-3 month-old) *Mef2c*-Het mice and their WT littermate controls for evaluation of hearing sensitivity. In agreement with a previous study in another mouse strain, ABR wave I threshold was lower in young adult C57BL/6J wild-type mice compared to animals at P16 (shortly after hearing onset) (Song et al., 2006) (**Fig. 2C,D**). *Mef2c*-Het mice showed threshold changes in young adults at 2-3 months. A significant elevation of ABR wave I thresholds was observed at most middle and low frequencies in young adult *Mef2c*-Het mice compared to their littermate wild type controls (**Fig. 2D**). Elevated thresholds ranged from 8 to 20 dB SPL (**Extended data Fig. 2-1**), suggesting a mild reduction of hearing sensitivity in young adult *Mef2c*-Het mice. No significant difference in ABR threshold was identified between male and female animals at age 2-3 months, suggesting sex is not an effect modifier for hearing sensitivity in *Mef2c*-Het mice (**Fig. 2E**).

**Figure 2.**
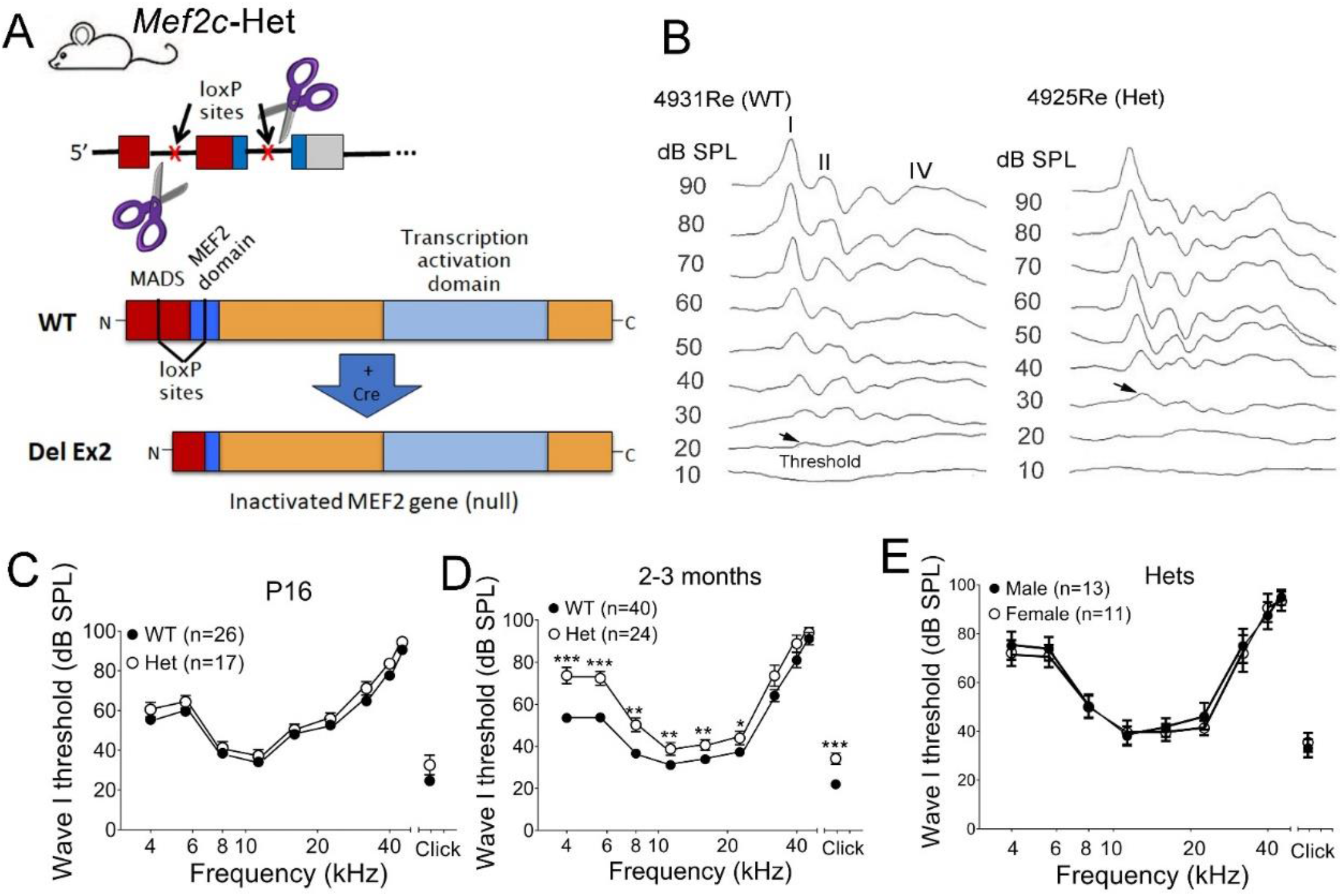
MEF2C deficiency causes a mild reduction of hearing sensitivity in young adult mice. ***A***, Schematic showing mouse *Mef2c* gene exons 1-3 and the approximate region eliminated (Exon 2; Ex2) in the global heterozygous *Mef2c* mutant (*Mef2c*-Het). Exon2 encodes a large part of the MADS/MEF2 domains (Harrington et al., 2020). ***B***, ABRs to click stimuli were recorded in a young adult *Mef2c*-Het mouse and a wild-type littermate control (WT). ABR waveforms from I to IV and an elevated ABR wave I threshold (arrows) in *Mef2c*-Het *mice* are shown for two representative young adult animals. ***C-E***, The averaged ABR wave I thresholds are shown in WTs and *Mef2c*-Het mice aged P16 (**C**), and 2-3 months (**D**). Significant threshold shifts were present in *Mef2c*-Het animals as compared to littermate controls (WTs) in both young adult groups (ABR wave I thresholds are presented as mean ±SEM, unpaired *t* test with Welch’s correction, \**p* <0.05; \*\**p*<0.01; \*\*\**p*<0.001) (detailed statistical analysis information was included in **Extended data Fig. 2-1**). ABR threshold difference was not identified between a group of *Mef2c*-Het mice and littermate controls (**E**).

### MEF2C deficiency results in reduced neural synchrony and auditory nerve activity

To further evaluate peripheral auditory function, we measured suprathreshold AN function using the phase-locking value (PLV). PLV is measured across single-trial ABR recording (Harris et al., 2018, 2021; McClaskey et al., 2020), and it quantifies the consistency of the phase of the response across individual trials (synchrony) within a given frequency range. The amplitude of the ABR is dependent on two independent factors: the number of neurons actively engaged by the stimulus and the synchrony of the response across trials (PLV). Factors that affect synchrony may therefore result in a smaller ABR amplitude despite a relatively preserved contingency of AN fibers. Similarly, a loss of AN fibers with preserved synchrony in the remaining fibers would result in a smaller peak amplitude. As shown in **Figures 3A** and **3B**, PLV was quantified at supra-threshold levels (60 dB SPL and above) and a linear mixed-effects regression was performed with stimulus level and genotype as fixed factors and mouse as a random factor, with stimulus level nested within the mouse. Stimulus level was a significant predictor of PLV [B = 0.014, t(70) = 9.861, p < .001], with PLV increasing with stimulus level for both genotypes. Most importantly, mouse genotype was a significant predictor of PLV [B = -0.375, t(75.76) = -3.241, *p* = 0.001], with PLV higher for WT mice than for *Mef2c*-Het mice. There was also a significant interaction between mouse genotype and stimulus level [B = .004, t(70) = 2.84, *p* = 0.006], indicating that the PLV of *Mef2c*-Het mice decreased with decreasing stimulus level to a stronger degree than for WT mice. In addition, post-hoc independent t-tests with Bonferroni correction showed that the difference between groups was significant at 60 dB SPL, where the PLV of *Mef2c*-Het mice (M = 0.337) was lower than the PLV of WT mice (M = 0.575) [t(10) = 3.402, *p* = 0.007, Cohen’s *d* = 2.152]. Together, these analyses revealed that Mef2c deficiency results in reduced neural synchrony at certain supra-threshold levels in the AN of young adult animals. To further characterize the AN dysfunction, we measured ABR wave I amplitude input/output functions (I/O; or amplitude growth functions), which allowed for evaluation of AN function at supra-threshold levels, in WT and *Mef2c*-Het mice (**Fig. 3C,D**). Linear mixed-effects regression revealed that stimulus level was a significant predictor of amplitude, with amplitude increasing with stimulus level for both genotypes. There was a significant interaction between mouse genotype and stimulus level at 8 kHz, demonstrating that amplitudes increase more with increasing in stimulus level in WTs compared to *Mef2c*-Het mice (see detailed information in **Extended Data in Fig. 3-2**) (**Fig. 3C**). In addition, the linear model analysis revealed significant effects of genotype on slope of the wave I amplitude growth function for both 8 and 11 kHz (**Fig. 3C,D**; Extended Data Fig. 3-2), indicating that AN activity was reduced at higher levels in *Mef2c*-Het mice. These observations further confirm the decline of suprathreshold AN activity in young adult *Mef2c*-Het mice.

**Figure 3.**
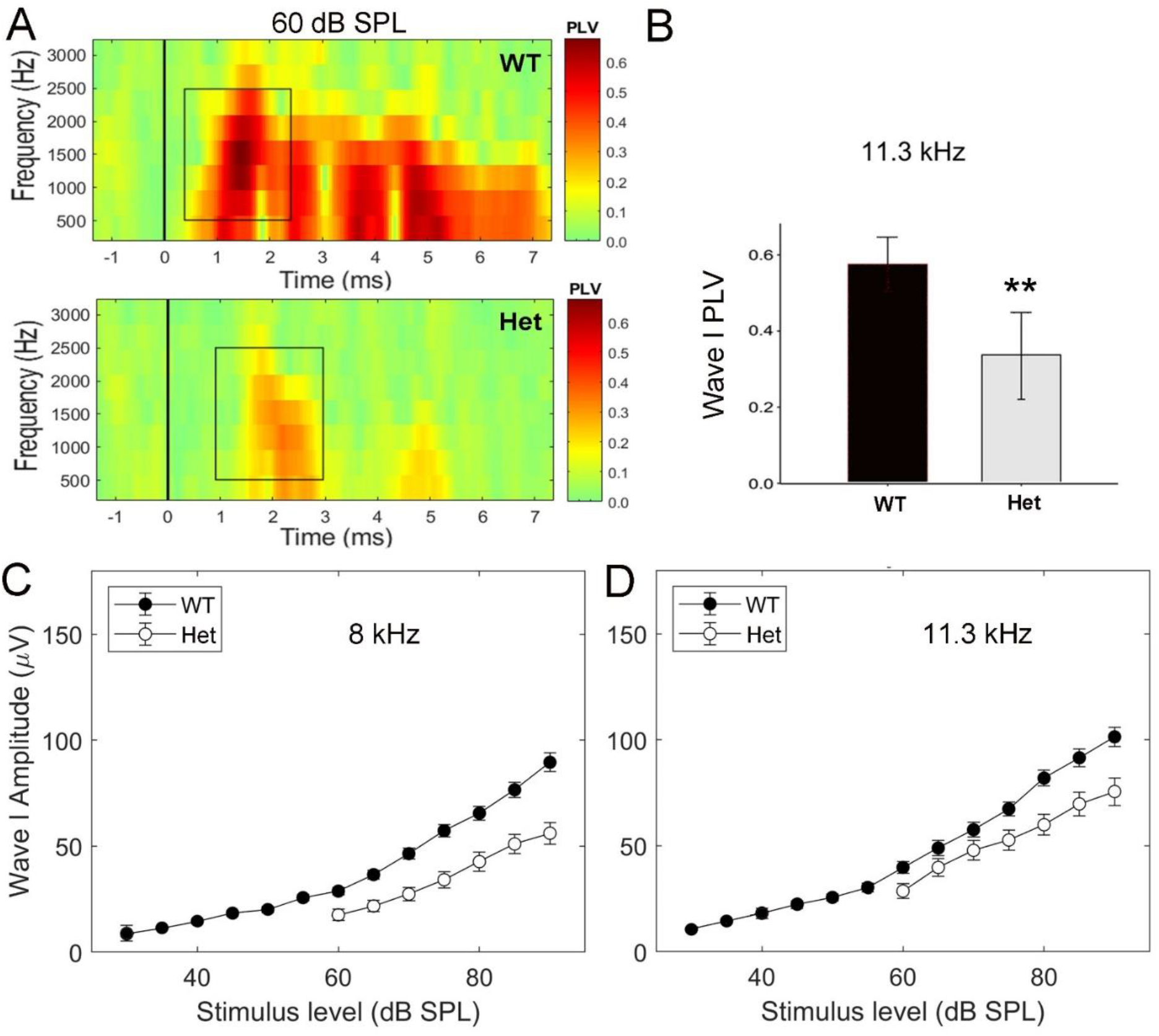
Reduced neural synchrony and AN activity in young adult *Mef2c*-Het mice. ***A***,***B***, Synchrony of AN firing of *Mef2c*-Het mice was poorer than that in WT mice. PLV reflects uniformity of phase at a specific time and frequency across trials and has been used to measure synchronization in neural activity from mouse ABR data. In the heat-map images in **A**, the X-axis represents time (signal onset = 1 ms); the y-axis represents frequency. Strength of PLV above baseline (green) is indicated by color (red). Independent t-tests with post-hoc Bonferroni corrections revealed that the difference between groups is most pronounced at 60 dB SPL, where the PLV of *Mef2c*-Het mice is significantly lower than the PLV of WT mice (n=6 animals per group; \*\**p*<0.01; detailed statistical information in **Extended data Fig. 3-1**). ***C***,***D***, ABR amplitude I/O functions at 8 and 11.3 kHz in WTs and *Mef2c*-Het mice. Note the reduced amplitudes of *Mef2c*-Het mice at high stimulus levels and a significant change in the slope of amplitude growth (from 75 to 90 dB SPL) when compared with WT, suggesting a decline of auditory suprathreshold function (*p* = 0.025 and 0.005 for 8 and 11.3 kHz, respectively; ABR wave I amplitudes are presented as mean ±SEM; n=24 animals for WTs and n=40 animals for Hets; detailed statistical information in **Extended data Fig. 3-2**).

### MEF2C deficiency leads to glial dysfunction in AN of young adult animals

To further elucidate the cellular and molecular changes that may contribute to these deficits, RNA-seq analysis was conducted on AN in *Mef2c*-Het mice at P21, a stage at which mouse ABR wave I threshold approximates the level of young adults (Song et al 2006). As shown in **Figure 4A-D**, RNA-seq analysis of WT and *Mef2c*-Het mice AN identified 258 genes that were differentially expressed (fold change >1.5 and *p*adj <0.10). Enrichment analysis of these genes showed that they were highly representative of neurogenic function, with the top 15 significant biological processes including categories relating to glia cells, myelination, and other neural developmental processes. To clarify these findings, all significant biological processes were refined by eliminating redundant categories. This refinement revealed that there was a cluster of highly similar processes that were centrally linked to sensory perception of sound. This cluster included several processes relating to glial cell function and myelination and also processes relating to the development of tissues derived from mesenchymal cell lineages.

**Figure 4.**
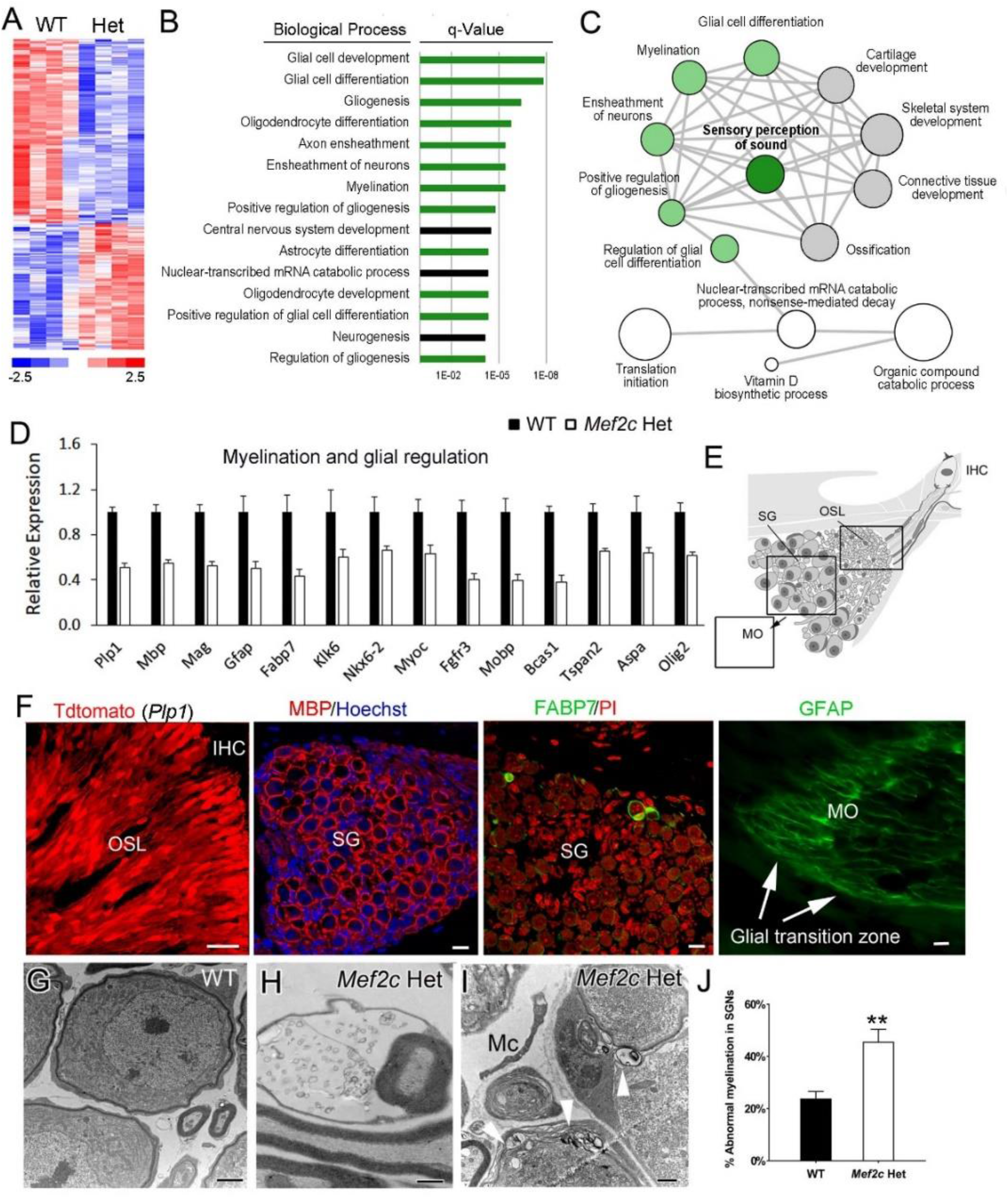
Gene expression alterations related to glial cell function and abnormal myelination in the AN of *Mef2c*-Het mice. ***A***, RNA-seq analysis identified 258 genes differentially expressed in AN of *Mef2c*-Het mice (**Extended Data Fig. 4-1**). Differential expression was defined as absolute fold change >1.5 and *p*adj (FDR step-up) <0.05 between P21 WT and *Mef2c*-Het mice (n=4 samples per group; 4 AN collected from 2 mice per sample). ***B***, Biological process enrichment results for differentially expressed genes shown in **A**. Significant gene terms and scores (Benjamini-Hochberg adjusted *p* <0.05) are shown. ***C***, Interactive graph of enriched biological process terms summarized with REVIGO Supek et al., 2011. Node size in **C** reflects the number of differentially expressed genes; colored nodes (green or gray) indicate the central grouping of highly similar categories relating to sensory perception of sound. Biological processes relating to glial cells are colored green in **B** and **C. *D***, Downregulation of genes related to myelination and glial regulation. ***E***, schematic image of AN cross-section illustrating areas along AN within osseous spiral lamina (OSL), spiral ganglion (SG) in Rosenthal’s canal, and modiolus (MO). ***F***, Immunohistochemical detection in mouse ANs for the glial cell markers that were found to be downregulated by RNA-seq. Shown are Schwan cells within OSL (*Plp1*), satellite cells around type I SGNs (MBP), satellite cells around type II SGNs (FABP7), and glial cells in the AN central process (GFAP) in MO. ***G***, EM graph showing healthy appearance of the SGNs in young adult WT animals. ***H-J***, Disruption of the myelin sheath surrounding an axon (**Fig. 4H**) and SGNs (white arrowheads; **Fig. 4I**). Percentage of SGNs with abnormal myelin sheaths was greatly increased in *Mef2c*-Het mice (Mann Whitney U test, *p*=0.0043, n=4 mice/group; **Fig. 4J**). Scale bars = 10 µm in **F**; 2 µm in **G**; 1 µm in **H**,**I**.

Proper glial cell development and function (e.g., myelination) is critical for AN structural and functional maturation (Panganiban et al., 2021). A small alteration in conduction velocity resulting from changes in myelin integrity may have significant effects on neuronal function such as spike-time arrival and neural synchrony (Pajevic et al., 2014). Here our RNA-seq analysis of *Mef2c*-Het mice found that there was downregulation of numerous glial cell markers, several of which have established importance in other nervous systems. These markers were detected in several different types of glial cells in AN of *Mef2c*-Het mice (**Fig. 4D-F**). For example, myelin proteolipid protein 1 (*Plp1*) was present in Schwann cells around AN axons within the osseous spiral ligament (OSL). Myelin basic protein (MBP) was located in satellite cells around SGNs and axons. Fatty Acid Binding Protein 7 (FABP7) was seen in satellite cells of a small subset of SGNs, type II SGNs, which are often found in the periphery of the spiral ganglion. Glial fibrillary acidic protein (GFAP) was expressed in the oligodendrocytes in the central AN process, which is located next to the glial transition zone (**Fig. 4F**). Ultrastructural examination of young adult AN myelin detected pathologies in *Mef2c*-Het mice (**Fig. 4G-J**). SGNs appeared normal in WT adult animals based on the neuronal cell bodies being completely enveloped by a thick, compact myelin sheath (**Fig. 4G**). In contrast, several features of degeneration were apparent in myelin sheaths of *Mef2c*-Het AN, including folded or collapsed lamella with balloon-like features (**Fig. 4H**), thinner lamellae, loose lamellae, or split myelin lamellae with discontinuities containing vacuole-like inclusions (**Fig.4I**) in glial cell cytoplasm. Disorganization of myelin was evident both in SGNs with severe degenerative changes and also in SGNs with a relatively normal appearance. Disrupted myelin sheaths were also seen in AN processes within Rosenthal’s canal and the osseous spiral lamina. Quantification of SGNs with myelin sheath abnormalities showed that occurrence was significantly increased in young adult *Mef2c*-Het mice compared to littermate WT controls. In 236 randomly selected SGNs from the middle turn sections of 4 WTs and 4 *Mef2c*-Het mice, 46±4% of SGNs in *Mef2c*-Het mice showed abnormal myelin compared to only 24±3 % in WT controls (*p*=0.002, Mann-Whitney U test; **Fig. 4J**).

### MEF2C deficiency causes aging-like alterations in neurons and mitochondria in young adult AN

Ultrastructural observations of *Mef2c*-Het AN detected evidence of SGN degeneration, including a loss of SGNs (**Fig. 5A**) and the presence of apoptotic bodies within dying SGNs (**Fig. 5B**). To quantify the effects on neuronal cell number, we performed immunolabeling studies in *Mef2c*-Het and littermate controls at both young adult (around 3 months) and older (around 6 months) stages to detect βIII-TUBULIN (TUBULIN), which has been previously shown to be highly expressed in type I SGNs of the young adult mouse (Sekerková et al., 2008; Jyothi et al., 2010). Interestingly, statistical analysis of the findings did not reveal a significant difference in numbers of neuronal cells between *Mef2c*-Het and littermate controls at either the young adult stage (**Fig. 5C**) (n=3 animals per group; p>0.05, Mann-Whitney U test) or older adult stage (**Extended data Fig. 5-2**) (*Mef2c*-Het, n=5; littermate WT control, n=7; *p*> 0.05, Mann-Whiney U test). This suggests that the number of affected cells in *Mef2-*Het mice may be a small fraction of the total. Consistent with these findings, count of CTBP2^+^ synapses of Type I SGNs revealed no significant difference in the synapse numbers at the apical-middle portion of the cochlear duct (**Fig. 5Q-T**).

**Figure 5.**
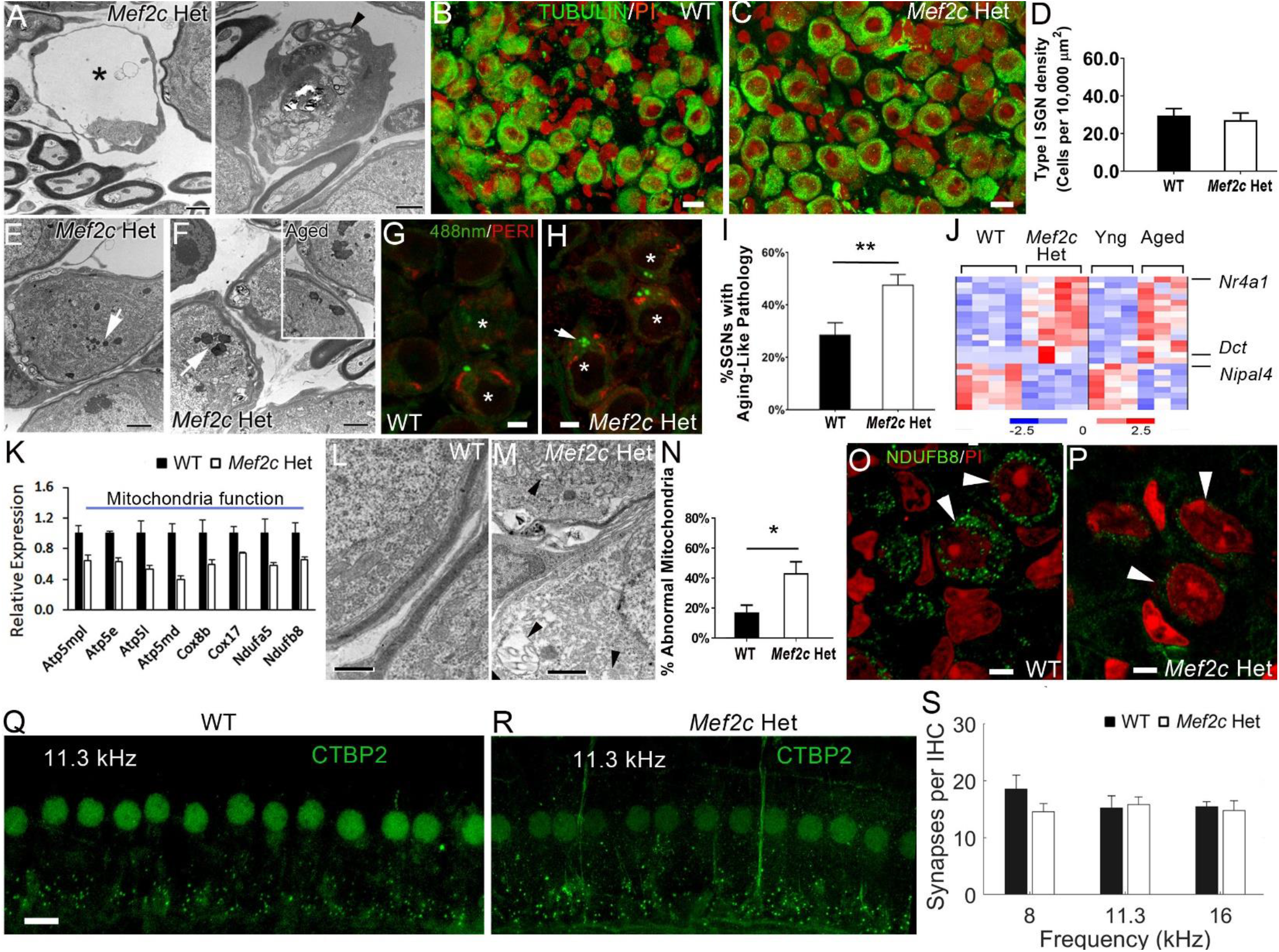
Aging-like neuronal changes and mitochondrial dysfunction in the ANs of *Mef2c*-Het mice. ***A***,***B***, Degenerative SGNs were seen in EM graphs of young adult *Mef2c*-Het mice. ***B-D***, No significant loss of SGNs was identified in young adult *Mef2c*-Het mice (*p>0*.*05*, Mann-Whitney U test; n=3 animals per group). Type I SGNs of young adult WT and *Mef2c*-Het mice were stained with anti-βIII-Tubulin (β-Tubulin; green) antibody. Nuclei were stained with propidium iodide (PI). ***E***,***F***,***I***, Increased aging-like pigment (white arrows in **E**,**F**) were seen in SGNs of young adult *Mef2c*-Het mice *(p=0*.*004*, Mann-Whitney U test; n=4 animals per group). The image in the right panel in **F** shows the aging pigment in SGNs of aged CBA/CaJ mice aged 2.5 years. ***G***,***H***, Aging pigment is detected by autofluorescence. Aging pigment was often seen in type I SGN soma that contained low intensity PERIPHERIN (PERI) labeled intermediate neurofilaments (red), but was not seen in areas in which nuclei were located (*). ***J***, Evaluation of RNA-seq data found that 28 genes differentially expressed in *Mef2c*-Het AN also changed in aging AN (gene list is included in **Extended Data Fig. 5-1**). ***K***, Mitochondrial function genes downregulated in *Mef2c*-Het AN. ***L-P***, *Mef2c*-Het mice mitochondrial abnormalities in AN. ***L***,***M*** TEM of young adult AN in WT and *Mef2c*-Het. Black arrowheads in **M** indicate abnormal mitochondria in SGNs. ***N***, *Mef2c*-Het mice display an increase number of abnormal mitochondria (*p*=0.029, Mann-Whitney U test, n=3 animals per group). ***O, P***, Altered distribution of the mitochondrial regulator NDUFB8 in SGN of *Mef2c*-Het AN. ***Q-S***, Number of ribbon synapses of type I SGNs in young adult *Mef2c*-Het mice appeared normal. Ribbon synapses, a subset of auditory nerve fibers and inner hair cell (IHC) nuclei were stained positively with anti-CTBP2 antibody. No significant change in CTBP2^+^ synapse number was seen in the 8, 11.3 and 16 kHz frequency areas of the *Mef2c*-Het mice (n=4-5) compared to WTs (n=3-5) (Type III ANOVA analysis; *p*=0.383, 0.705, and 0.406 for main effects of genotype, frequency, and the interaction of genotype*frequency, respectively). Scale bars = 2 µm in **A**,**E**,**F**; 10 µm in **B**,**C**,**Q**; 5 µm in **G**,**H**,**O**,**P**; 1 µm in **L**,**M**.

One of the striking hallmarks in aging neuronal cells is the accumulation of lipofuscin granules or neuromelanin pigments (also named “aging pigments”) in the cellular compartment of the soma as shown in SGNs of an aged CBA/CaJ mouse (**Left panel of Fig. 5F**; Riga et al., 2006; Sulzer et al., 2008; Jung et al., 2010). Interestingly, aging-like SGNs were more often seen in young adult *Mef2c*-Het mice than their WT littermate controls (**Fig. 5E; Right panel of Fig. 5F**). As shown in **Figure 5I**, quantitative analysis of EM graphs from middle turns of young adult mice revealed that 47±4% of SGNs in *Mef2c*-Het mice had at least two lipofuscin granules while only 29±5 % of SGNs in WT had this aging-like characteristic. The increase in these lipofuscin (or neuromelanin) inclusions was also validated by fluorescence microscopic detection of the autofluorescent inclusions (**Fig. 5G,H;** Jung et al., 2010). To further evaluate the premature aging-like phenomenon in young adult *Mef2c*-Het mice SGNs, we examined expression profiles of DEGs obtained from our RNA-seq analysis of *Mef2c*-Het AN using a prior RNA-seq dataset that tested the effect of aging (Panganiban et al., 2021). Our analysis found that 28 of the genes affected in *Mef2c*-Het AN were also significantly affected (*p*adj<0.1) in aging AN and agreed in direction of change (**Fig. 5J**). Dopachrome tautomerase (*Dct*) is a type I membrane protein and a key regulator of biosynthesis of melanin, including neuromelanin, which can form a similar type of pigment accumulation in aging nervous tissue (Costin et al., 2005; Double et al., 2008). Nipa-like domain-containing 4 (*Nipal4*) is involved in synthesis of long-chain fatty acids and plays a role in lipid metabolism (Dahlqvist et al., 2012). Nuclear receptor 4A1 (*Nr4a1*) is a mediator of macrophage function and plays a role in regulating inflammatory responses. Translocation of Nr4a1 protein from the nucleus to mitochondria leads to apoptosis (Bouzas-Rodríguez et al., 2012). Taken together, combined quantitative ultrastructural evaluation and RNA-seq analysis revealed subcellular and molecular evidence of AN dysfunction in young adult *Mef2c*-Het mice.

Ultrastructural examination of young adult mouse AN found that there were pathological alterations in some mitochondria and that these alterations were more frequently seen in *Mef2c*-Het animals (**Fig. 5L-N**). Alterations included increased intra-organelle space and separation of cristae. A count of healthy and pathological mitochondria was completed on randomly selected axon EM graphs that were taken from the middle portions of cochlea from WT and *Mef2c*-Het mice. Our data revealed a significant increase in abnormal mitochondria in the *Mef2c*-Het mice. Approximately 43% of examined mitochondria were abnormal in *Mef2c*-Het mice (n=4); whereas, only 17% of examined mitochondria were abnormal in WT controls. To further validate these results, a re-examination of RNA-seq data for the 258 DEGs affected in *Mef2c*-Het AN found that eight genes were related to mitochondria function and all were downregulated (**Fig. 5K**). Immunohistochemical analysis for one of these, NADH dehydrogenase [ubiquinone] 1 beta subcomplex \subunit 8 (NDUFB8), found that expression of Ndufb8 was high in mitochondria of SGNs in WT mice, but it was weak in SGN of *Mef2c*-Het mice (**Fig. 5K,O,P**). Again, quantitative ultrastructural evaluation and gene expression analysis revealed that mitochondrial dysfunction may be one mechanism underlying AN functional decline in the *Mef2c*-Het mice.

### *Mef2c*-Het mice exhibit increased inflammation and macrophage activation in AN and stria vascularis

Our previous study showed that Mef2c hypofunction in microglia leads to dysregulation of microglial genes and ASD-related behaviors (Harrington et al., 2020). Based on those findings, we hypothesized that Mef2c deficiency would cause abnormal macrophage activity in the AN and its surrounding cochlear environment (e.g., cochlear vasculature in the AN and stria vascularis). Cochlear ultrastructural examination (**Figs. 6A-D, F.G**), gene expression analysis of AN (**Fig. 6E**), and immunohistochemical evaluation of AN macrophages and microvasculature (**Fig.6H**) and stria vascularis (**Fig. 6I,J**) all yielded results that support this hypothesis. First, microvasculature was disrupted in young adult *Mef2c*-Het AN (**Fig.6B,C**) and stria vascularis (**Fig.6D**), and macrophages can be seen interacting with the abnormal vasculature. Second, RNA-seq findings showed significant upregulation of anti-angiogenic genes and downregulation of pro-angiogenic genes in *Mef2c*-Het AN (**Extended Data Figure 6-1**). Third, RNA-seq findings showed upregulation of genes relating to chronic inflammation and macrophage activation, but downregulation of genes involved with innate immune response and anti-inflammation in *Mef2c*-Het AN (**Fig. 6E**), including *S100a8, S100a9, Chil3*, and *Cd177*. Qualitative and quantitative morphometric analysis further showed that macrophage activation was increased (**Fig.6F-J**) and macrophage and microvasculature interactions affected in *Mef2c*-Het mice (**Fig.6I**). TEM revealed interactions between macrophages and neurons and the uptake of myelin debris through phagocytosis (**Fig. 6F**). The presence of myelin-filled lysosomes within macrophages is suggestive of demyelination and neuron degeneration, as previously observed in a noise-induced AN degeneration model (Panganniban et al., 2018). Immunohistochemistry performed on 20 µm-thick AN sections and whole-mount preparations of stria vascularis show that there was an increase in the number of macrophages with activated morphology, especially in the apical turn, based on retracted processes and reduced cellular area and volume (**Fig. 6H-J;** p<0.0001 (for both cellular area and volume measure), unpaired, two tailed t test with Welch’s correction). To further test the impact of MEF2C deficiency on the strial function, we quantitatively evaluated the immunoactivity inwardly rectifying potassium (Kir) channel 4.1 (Kir4.1) (**Extended data Fig.6-2**). Kir 4.1 is 616 expressed in intermediate cells and plays an important role in the maintenance of the high K^+^ concentration and generation of endocochlear potential (Liu et al., 2019). Our data revealed there are no significant changes in Kir4.1 immunoreactivity of the strial intermediate cells of *Mef2c* Hets compared to that of WTs, suggesting limited impact of MEF2C deficiency in cochlear lateral function and EP level.

**Figure 6.**
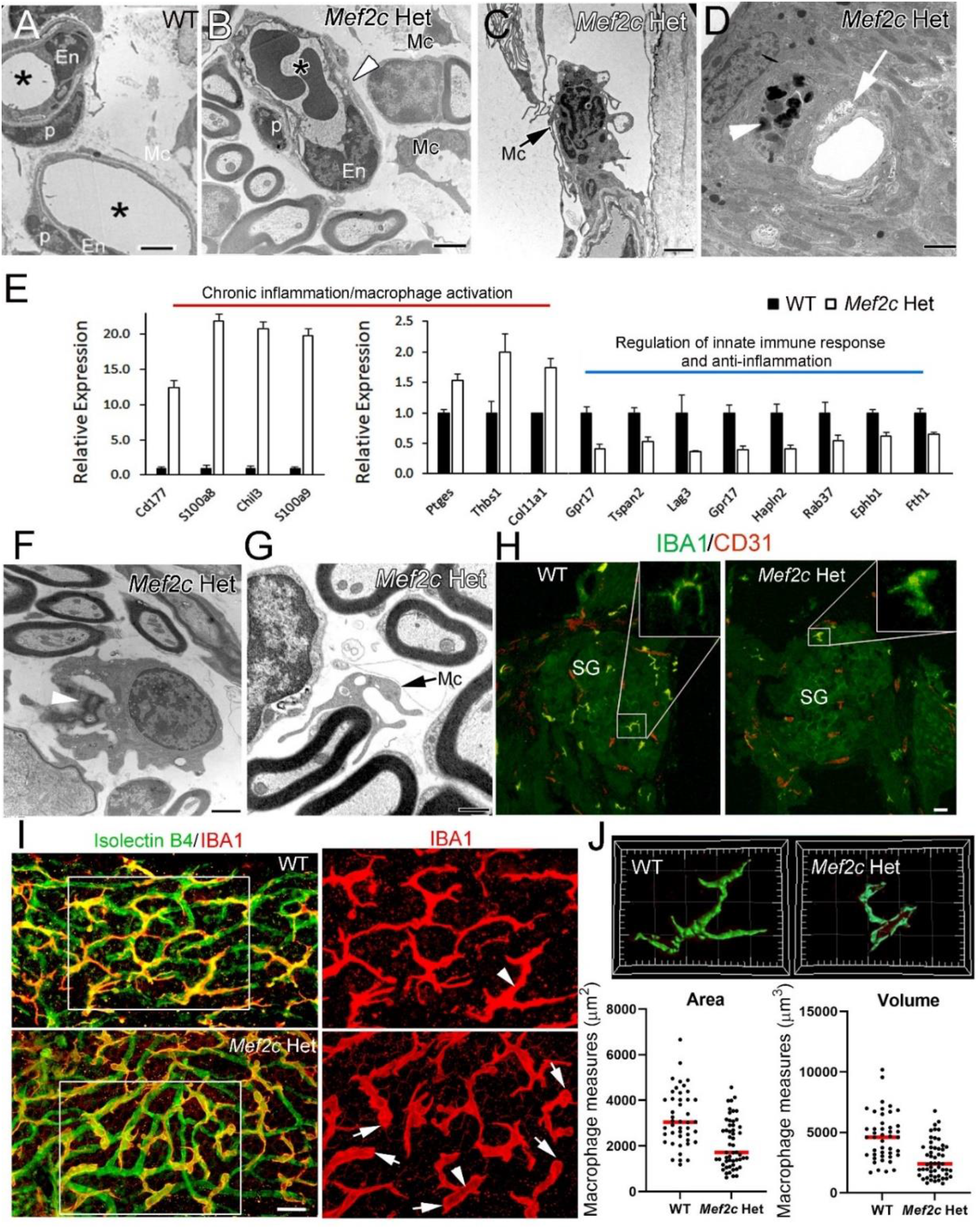
Increased cochlear inflammation and macrophage activation in AN and stria vascularis of young adult *Mef2c*-Het mice. ***A-D***, EM graphs show pathological alteration in microvasculature (white arrowhead in **B**; white arrow in **D**), frequently engaging with activated macrophage (Mc, black arrow in **C**), in *Mef2c*-Het AN (**B**,**C**), and stria vascularis (**D**). TEM graph in A shows endothelial cells (En) adjacent to pericytes (p) in a normal-appearing microvasculature in AN. Asterisks indicate capillary lumens. A white arrowhead in **D** indicates a perivascular macrophage-like melanocyte (characterization based on previous studies by Zhang et al., 2012) located around the disrupted microvasculature in stria vascularis of a *Mef2c*-Het mouse. ***E***, Changes in expression of genes related to chronic inflammation/macrophage activation, regulation of innate immune response and anti-inflammation. ***F***,***G***, TEM graphs show a macrophage cell engulfing myelin debris (**F**) and a macrophage process surrounding a healthy axon (**G**). ***H-J***, Increase in macrophages with activated morphology and changes in interaction between macrophages and microvasculature in AN and stria vascularis of young adult *Mef2c*-Het mice. CD31 antibody (**H**) or Isolectin GS-IB4 conjugated with Alexa FluorT^M^ (**I**) were used for labeling microvasculature. IBA1 antibody was used as a marker for identifying macrophages. Images in Bottom panel show 3D views of selected macrophages in **I** (white arrow heads) generated with the Surface module of IMARIS (IMARISx64 9.3.1). Quantitative analysis of macrophage morphology reveals reduced surface area and cellular volume in the stria vascularis of young adult *Mef2c*-Het mice (Bottom panel in **J**; 44 and 53 macrophages were randomly selected and measured from 3 WTs and 3 Hets, respectively; the detailed statistical analysis information in **Extended data Fig. 6-1**). Scale bars = 2 µm in **A-D**,**F**; 800nm in **G**; 15 µm in **H**; 20 µm in **I**.

To further test the hypothesis that Mef2c deficiency in immune cells (such as Cx3cr1 expressing cochlear macrophage) leads to cochlear pathology and AN functional decline, young adult *Mef2c*^*flox/+*^; *Cx3cr1-Cre* mice were examined for ABR wave I threshold and supra-threshold function. As shown in **Extended data Fig. 6-3** and **6-4**, there were no significant alterations in wave I ABR threshold or supra-threshold function in *Mef2c*^*flox/+*^; *Cx3cr1-Cre* mice as compared to that of their littermate controls, suggesting that deficiency of Mef2c in immune cells alone may not lead to AN functional decline.

### Mef2c deficiency has no gross effect on hair cell function and cochlear bone formation

Since *Mef2c*-Het mice showed a significant reduction of ABR wave I threshold in low-and middle-frequency ranges compared to controls, we examined whether there were functional declines or pathological alterations in sensory hair cells. Functional and structural analyses of sensory hair cells in *Mef2c*-Het mice did not detect an overt effect on sensory hair cells (**Figure 7**). That was supported by cochlear microphonic (CM) measurement which detected no significant difference between the young adult WTs and *Mef2c*-Het mice (**Fig. 7A**), suggesting no significant loss of outer hair cell function in *Mef2c*-Het mice. Furthermore, histological analysis did not reveal gross morphological changes in apical and middle portion cochlear structure, using either cochlear sections (**B**) or whole-mount preparations (**C, D**). Since MEF2C protein was detected in bone forming cells around the cochlear duct (**Extended data Fig. 1-3**), we next examined whether MEF2C deficiency affected bone formationin the ossicles and cochlear bony capsule. As shown in **Figure 7G-H**, no gross changes were evident in the cochlear bony capsule (**D**), ossicles (**H**) or bone structure around the AN (**I**), suggesting that MEF2C deficiency was not sufficienct to disrupt middle and innea ear bone formation

**Figure 7.**
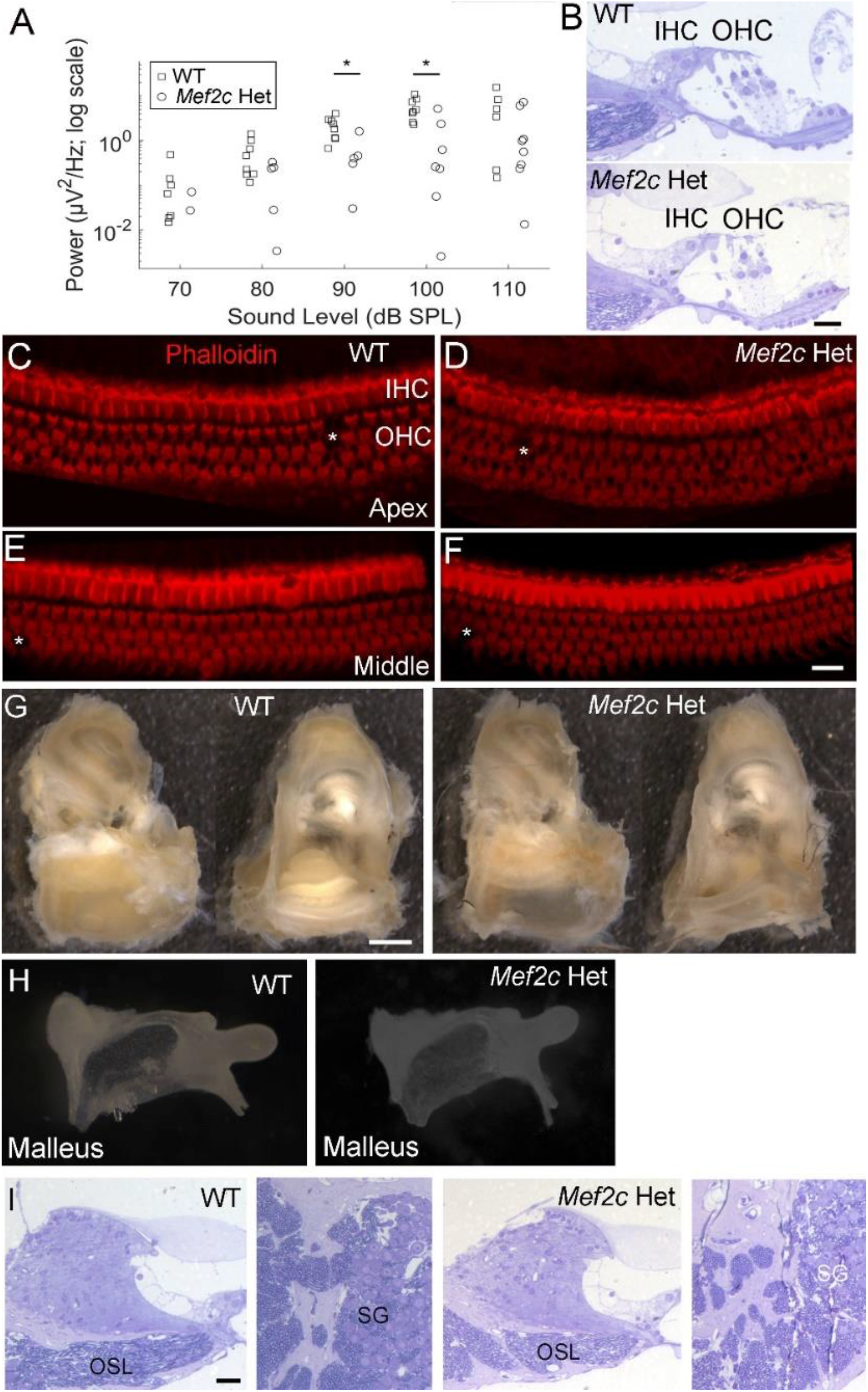
Evaluation of sensory hair cells, ossicles and cochlear bony capsules in young adult *Mef2c*-Het mice. ***A***, No difference in CM response was identified between young adult WT and *Mef2c*-Het mice (**Extended data Fig. 7-1** for detailed statistical analysis information). ***B***, Representative images of toluidine blue-stained organs of Corti from the middle portion of the cochlea showing IHCs and outer hair cells (OHC) in young adult WT and *Mef2c*-Het mice. ***C-F***, Surface preparations from apical (**C**,**E**) and middle (**D**,**F**) portions of cochlea from WT (**C**,**E**) and *Mef2c*-Het (**D**,**F**) cochleae. Very infrequent hair cells loss (*) was seen in both groups. ***G***,***H***, No significant change was observed in the bony inner ear portion of temporal bones (**G**; Left side image shows side view of cochlea; right side shows view from the internal auditory canal) or ossicles (**H**) of young adult *Mef2c*-Het mice. ***I***, Representative images of toluidine blue-stained AN from the middle portion of young adult WT and *Mef2c*-Het mice showing the bony structures around OSL and SG. Scale bars = 20 µm in **B-F**,**I**; 600 µm in **G**.

## Discussion

Establishment of animal models of human ASD with well-characterized AN functional deficiency is a crucial first step toward understanding the contribution of peripheral AN dysfunction to communication deficits and other ASD behaviors. However, to date only a small number of auditory investigations have been done with ASD animal models. Among those studies, peripheral AN functional decline was well-characterized in a mouse model of Fragile X syndrome (*Fmr*1 knockout mice) (Rotschafer et al., 2015). ABR wave I amplitude growth function measures in *Fmr1*^*-/-*^ mice suggested a decline in peripheral AN activity, together with an approximate 10 dB SPL threshold elevation and deficits in auditory brainstem function. Previous studies also showed that mouse and rat models of Fragile X syndrome exhibit hypersensitivity and auditory processing dysfunction (Rotschafer and Razak 2013; Garcia-Pino et al., 2017; McCullagh et al., 2020; El-Hassar et al., 2019; Auerbach et al 2021). In addition, abnormal ABRs were reported in *Adnp* haploinsufficient (*Adnp*^+/-^) mice, another animal model of ASD-linked gene deficiency (Hacohen-Keiman et al., 2019). Those studies revealed that ASD models exhibited auditory impairment but provided only limited information regarding possible changes in peripheral auditory organ or pathological alterations of AN degeneration and dysfunction, which is critical for understanding the link between AN function and ASD characteristics, such as sensory hypersensitivity and social communication deficits. Here we report that a model of human MEF2C hypofunction with well-defined ASD-like behaviors (Harrington et a., 2020) has a modest reduction in hearing sensitivity combined with AN functional deficits. This study is significant in that it: 1) links a monogenic ASD animal model with a clear impairment in peripheral AN structure and function, and 2) reveals a critical new role for Mef2c function in normal peripheral AN development and function.

Functional and structural integrity of the peripheral AN is critical for proper auditory processing in the central auditory system. However, ABR wave I threshold only represents the function of the most sensitive AN fibers; therefore, threshold is not regarded as a sensitive approach for assessing AN integrity. For example, a well-established mouse model of noise-induced hearing loss showed robust primary AN degeneration, but only a limited change in the threshold of ABR wave I (or the AN compound action potential threshold) (Kujawa and Liberman 2009; Liberman 2017). Measures of AN threshold are dependent on a small subset of AN fibers with high SR and low thresholds. In contrast, suprathreshold measures of AN function may reflect the contribution of AN fibers with varied SR and thresholds depending on level and stimulus parameters of the eliciting stimulus. Combining *in vivo* non-invasive measurement of AN synchrony by PLV together with ABR wave I amplitude growth functions (AN suprathreshold measures), our study found that *Mef2c*-Het mice have suprathreshold peripheral AN functional decline. For example, at 11.3 kHz, the frequency showing AN functional decline, the elevated thresholds were less than 9 dB SPL, with no significant threshold shifts in the higher frequencies (**Fig. 2D; Extended data Fig. 2-1**). These results agree with previous findings that there was no association between hearing loss and human MCHS, although difficulty with speech, intellectual disabilities, seizures, stereotypic movements, vision issues, and other ASD-like behaviors were well-documented in MCHS patients (Cooley Coleman et al., 2021). While there remains only limited evidence in humans that AN dysfunction is linked to ASD (Santos et al., 2017), peripheral AN deficits are known to contribute to changes in higher levels of the central auditory system, including sensory hypersensitivity and auditory processing dysfunction. Our recent studies reported that poorer PLV in humans with otherwise normal audiograms is associated with poorer auditory processing and speech recognition (Harris et al., 2017; 2021). Taken together, results of the current study highlight the potential clinical utility of suprathreshold peripheral AN functional evaluation as one potential diagnostic indicator of ASD. This technique was first developed and tested in human subjects, data can be acquired without active subject participation, and the procedure is relatively non-invasive (Harris et al., 2017; 2021).

Another important finding of this study is that the *Mef2c*-Het mice exhibit multiple cellular and molecular alterations that likely contribute to the peripheral AN functional decline. These alterations included glial cell dysfunction and myelin abnormality, aging-like SGN degeneration (e.g., lipofuscin and/or neuromelanin accumulation), neuronal mitochondrial dysfunction, and increased macrophage activation and inflammation. A microdeletion or point mutation within the protein-coding region of the *MEF2C* gene on chromosome 5 results in reduced MEF2C protein levels and function is one established cause of human MCHS (LeMeur et al., 2010; Mikhail et al., 2011; Tonk et la., 2011). Brain MRI studies in MCHS patients suggested that MEF2C mutation is also associated with delayed neuron myelination (Zweier et al., 2010; Rocha et al 2016). In agreement with this observation, our analysis of *Mef2c*-Het RNA-seq and ultrastructural data support the finding of glia dysfunction and abnormal myelination. Interestingly, while MEF2C was found to be present in many different cell types within adult AN, it was not found in mature glial cells (**Fig. 1**). This suggests that the observed consequences on glial cell gene expression and myelination are not due to MEF2C function within mature glia but may be the result of signaling from other cell types or as a secondary response to SGN dysfunction and/or increased cochlear inflammation and macrophage activation. Bioinformatic analysis of genes identified as ASD-related and AN developmentally regulated found that 18 were also putative MEF2C targets in microglia (Deczkowska et al., 2017; Bjorness et al 2020). Interestingly, the molecular function “signaling receptor binding” was among the most enriched category for these genes, suggesting the potential for MEF2C to elicit an effect on glia through a neuron-to-glia signaling mechanism. Notable among the genes was acetylcholinesterase (Ache), which is a chemical transmitter that regulates neuron-glial crosstalk and neuropathic pain in sensory ganglion (Matsuka et al., 2019).

Although no significant neuronal cell loss was found in *Mef2c*-Het mice, AN functional decline was supported by aging-like SGN degeneration including neuronal mitochondrial dysfunction. Previous studies revealed that MEF2 family members regulate transcription of both the nuclear and mitochondrial genomes and that their inhibition can cause decreased activity of mitochondrial complex I (ubiquinone oxidoreductase), increased hydrogen peroxidase levels, and reduced ATP production (Naya et al., 2002; She et al., 2011). In skeletal muscle, MEF2C plays a role in regulating metabolic homeostasis and affects animal body size (Anderson et al., 2015). The importance of the link between MEF2 and mitochondrial function was also highlighted in investigations of neuronal structural and functional plasticity (Brusco and Haas 2015). In the AN of young adult *Mef2c*-Het mice, our RNA-seq analysis detected downregulation of several mitochondrial genes, including mitochondrial complex I protein NDUFB8 (NADH dehydrogenase [ubiquinone] 1 beta subcomplex subunit 8). Distribution of NDUFB8 protein was also found to be affected in neuronal soma of *Mef2c*-Het AN. Together with the observed disruption of mitochondrial structure, the dysregulation of mitochondrial genes related to complex I activity suggests that mitochondrial hypofunction may be a cause of AN functional decline in *Mef2c*-Het mice. Previous studies showed that reduced mitochondrial complex I activity contributes to mitophagy dysfunction, which results in an increased number of damaged mitochondria in neuronal cells in neurodegenerative diseases (Franco-Iborra, et al., 2018). Accumulation of damaged mitochondria and autophagic vacuoles filled with lipofuscin or neuromelanin pigments are hallmarks of age-related neurodegeneration and are also associated with neuronal loss (Sulzer et al., 2008; Fang et al., 2019; Moreno-Garcia et al., 2018). Previous studies also suggest that an interaction between senescent lipofuscin and mitochondria may play a role in progression of cellular degeneration (Terman et al., 2007; König et al., 2017). Based on these observations, we hypothesize that *Mef2c* hypofunction contributes to aging-like mouse AN degeneration via a negative impact on mitochondrial function and cellular metabolism.

Transcriptomic analysis of postmortem brain tissues from individuals with autism has revealed expression changes in microglial-specific genes and genes related to inflammatory response (Voineagu et al., 2011; Gupta et al., 2014; Velmeshev et al., 2019). Recent studies revealed that MEF2C expression in microglia/macrophages is also important for brain development and maintenance of cognitive function in adults (Harrington et al., 2020; Deczkowska et al., 2017). Here our study revealed that MEF2C is expressed in neurons as well as in microglia/macrophages in the AN of the peripheral auditory system. In the peripheral auditory system, we found that *Mef2c*-Het mice demonstrated an increase in macrophage activation in the AN and the stria vascularis, an increase in mRNA expression of pro-inflammatory genes, and morphological disruption of microvasculature. There are at least three factors that may lead to increased IBA1 expressing cells in the cochlea: 1) as a direct result of Mef2c deficiency (since MEF2C is highly expressed in IBA1^+^ cochlear macrophage); 2) as a result of dysfunction of the spiral ganglion neuron, and 3) a consequence of the degeneration of glial cells in the AN. Our data showing increased inflammatory cells in the strial vascularis also suggest the direct impact of Mef2c deficiency in cochlear macrophages in multiple cochlear locations. Though the current study did not establish whether increased macrophage activation was a direct result of *Mef2c* hypofunction in those cells, SGN aging-like degeneration, or both, our data clearly reveal that the AN functional decline is accompanied by increased macrophage activation and a shift in the cochlea toward pro-inflammatory status.

Conditional deletion of one copy of *Mef2c* in *Cx3Cr1*-expressing cells (macrophages and other immune cell subtypes) in mice [*Mef2c*^*flox/+*^; *Cx3cr1-Cre* (*Mef2c cHet*^*Cx3cr1*^) mice] resulted in social interaction deficits and increased repetitive behavior (Harrington et al., 2020), suggesting that *MEF2C* haploinsufficiency in neuroimmune cells may produce ASD-related behaviors. However, in the peripheral auditory system, no significant alterations in auditory function were observed in the *Mef2c cHet*^*Cx3cr1*^ mice, suggesting a partial deletion of *Mef2c* in cochlear macrophages alone may not be enough to cause auditory nerve functional declines. Further study is needed to address the specific mechanistic relationships and whether inhibition of macrophage activation is a potential treatment strategy for improving AN function and reducing other ASD-related communication impairments.

Finally, our study provides a paradigm for evaluating AN functional decline at the physiological, cellular, and molecular levels in mouse models with only minimal hearing threshold changes. Comprehensive structural and functional characterization of the peripheral AN is a critical step in evaluating the relationship between peripheral auditory function and sensory hypersensitivity and other auditory processing impairments, whether in ASD or other common neurodevelopmental and neurodegenerative disorders. Measurement of ABR threshold and standard histological evaluation such as SGN counting are not sensitive approaches for identifying AN functional decline with mild or no hearing threshold elevation (termed “hidden hearing loss”).

Taking advantage of high-resolution imaging technology, the establishment of cochlear synaptopathy (loss of synaptic connections between AN fibers and cochlear hair cells) demonstrates the importance of primary AN degeneration in several forms of sensorineural hearing loss (Liberman and Kujawa 2017; Liberman 2017). The paradigm presented here combines comprehensive AN physiology (including inter-trial coherence) with high-resolution imaging of non-neuronal cells and transcriptomic analysis for evaluating the functional state of the peripheral AN. Using a model that displays a mild level of hearing loss, the application of this approach was able to detect deficiency in several non-neuronal elements, including glial cell dysfunction, myelination abnormality, alteration in cochlear microvasculature, and elevated macrophage activation that may directly or indirectly contribute to AN functional declines and degeneration.

## Acknowledgments

This work was supported by National Institutes of Health Grants K18DC018517 (H.L.), R01DC012058 (H.L.), P50DC000422 (H.L., K.C.H.), SFARI Pilot Award #649452 (C.W.C, H.L), R01 MH111464 (C.W.C.), T32014435 (C.M.M), P30GM103342 (J.L.B.), P20GM103499 (J.L.B.), and P30CA138313 from South Carolina Translational Science Laboratory Shared Resources, Hollings Cancer Center at the Medical University of South Carolina (MUSC). The Cell & Molecular Imaging Shared Resource was also supported by the SC COBRE in Oxidants, Redox Balance, and Stress Signaling (P20 GM103542), the SC COBRE in Digestive and Liver Diseases (P20 GM130457), the MUSC Digestive Disease Core Center (P30 DK123704) and the Shared Instrumentation Grants S10 OD018113. Bioinformatic analysis of RNA-seq data was done through the MUSC Proteogenomics Facility that is supported by NIGMS (GM103499) and the MUSC Office of the Vice President for Research. We thank John Lemasters for his suggestions with the use of the mitochondrial functional markers. We thank Catherine Bridges, Cindy Wang, Shelby Storm, Juhong Zhu, Carlene Brandon, Jiaying Wu, Logan Martin and Nancy Smythe for their excellent technical assistance, and thank Richard Schmiedt for his help with the system setup of single-trial ABR recording. We also thank Jayne Ahlstrom, Nathaniel Parsons and Cindy Wang for their comments and editing of the manuscript.

---

**Extended data Fig. 1-3 (Images).**
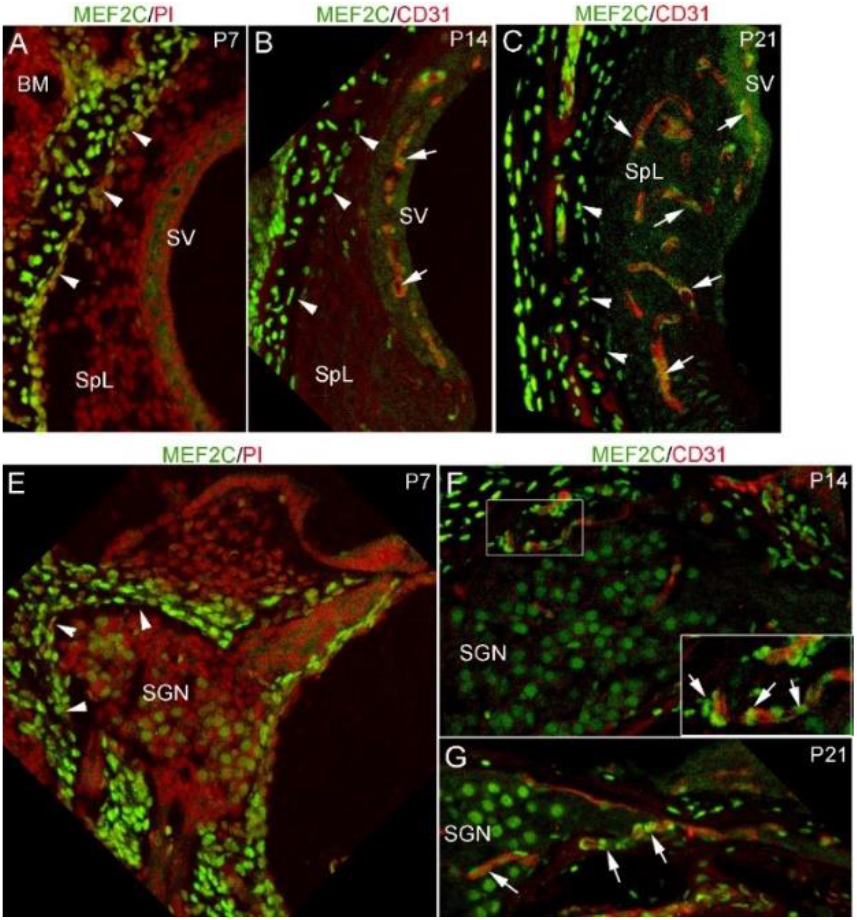
***A-G***, MEF2C expressing cells present around a blood vessel in AN (arrows in **F)** and cochlear lateral wall (arrows in **B**) and areas near bone-forming cells (arrowheads in **B**,**C**, and **E**) in postnatal developing cochlea.

**Extended data Fig. 5-2 (Images).**
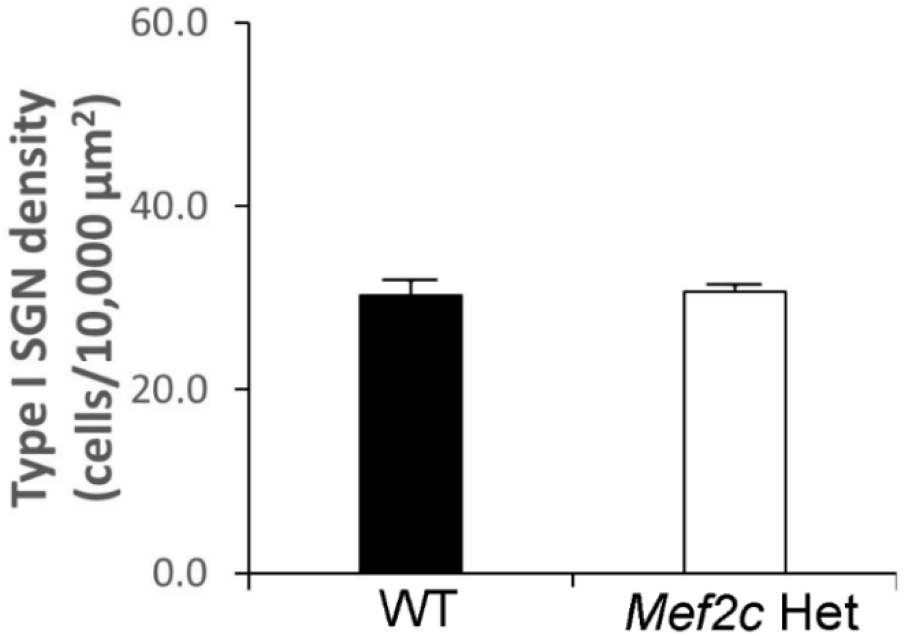
No significant loss of SGNs was identified in adult *Mef2c*-Het mice aged 6-7 months. No significant loss of SGNs was identified in *Mef2c*-Het mice (*p=0*.*5*, Mann-Whitney U test; n=7 for WTs and n=5 for *Mef2c* Hets). Type I SGNs of young adult WT and *Mef2c*-Het mice were stained with anti-βIII-Tubulin antibody.

**Extended data Fig. 6-2 (Images).**
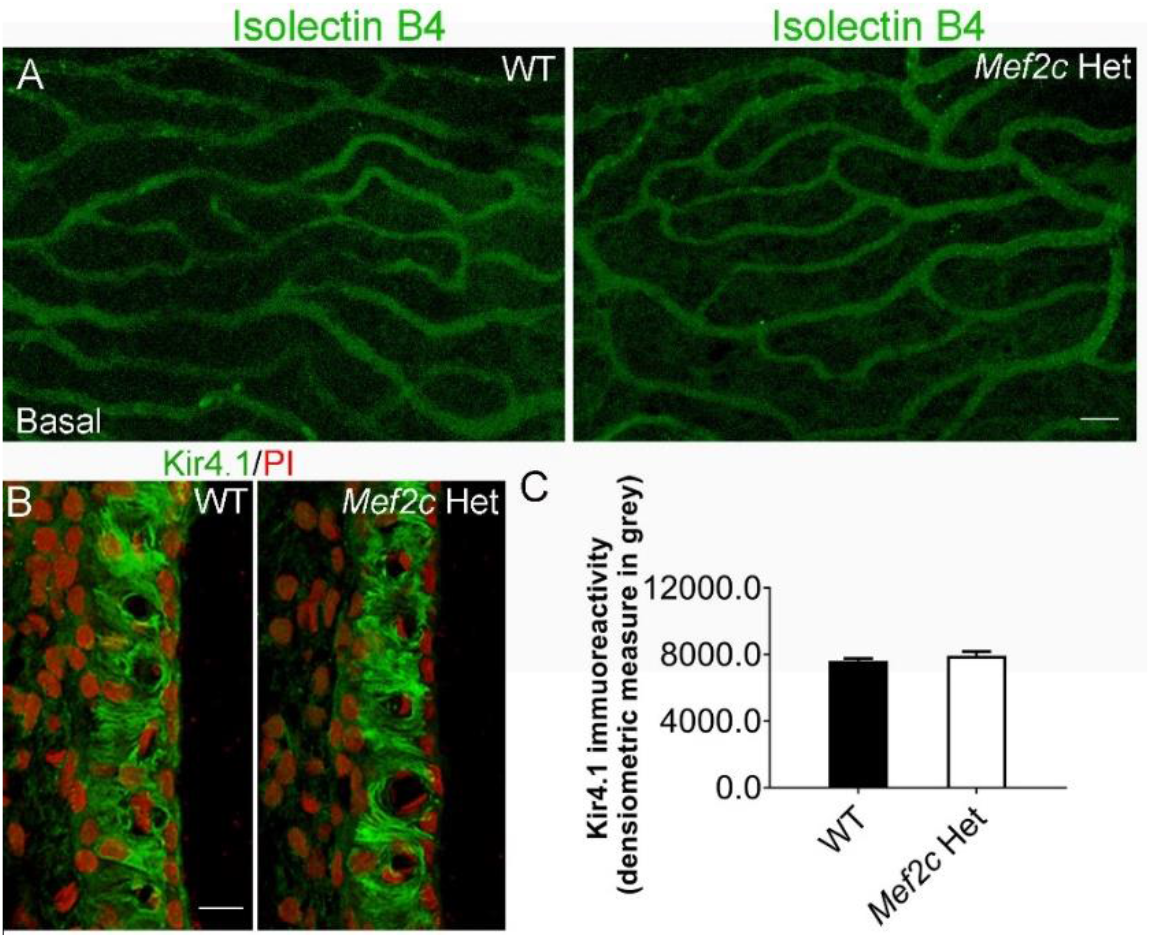
***A***, The strial microvasculature was stained with isolectin GS-IB4 conjugated with Alexa FluorT^M^ for young adult WT and Mef2c Het mice. ***B***, No significant change in Kir4.1 immunoactivity was identified in the strial vascularis of young adult Mef2c Het animals compared to that of WTs (*p=0*.*3*, Mann-Whitney U test; n=3 for WT and n=4 for *Mef2c* Het).

**Extended data Fig. 6-3 (Images).**
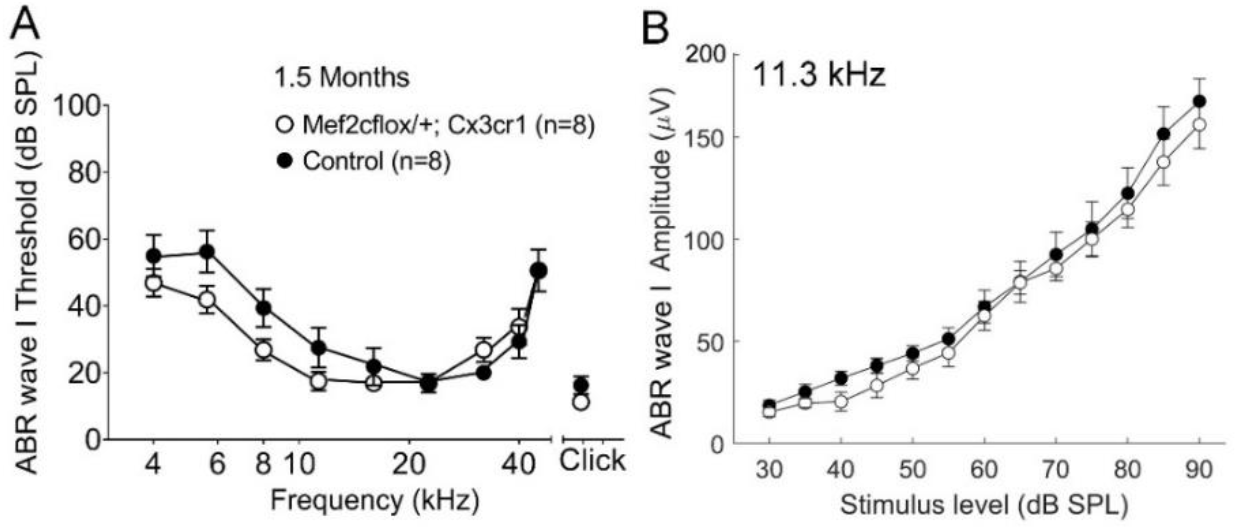
***A***, The averaged ABR wave I thresholds are shown in young adult WTs and *Mef2c cHet*^*Cx3cr1*^mice. No significant threshold shifts were seen in *Mef2c cHet*^*Cx3cr1*^ animals as compared to littermate controls (WTs). ***B***, No significant change was seen in ABR wave I amplitude I/O functions at 11.3 kHz between WTs and *Mef2c cHet*^*Cx3cr1*^ mice.

